# An evolutionary-conserved VPS34-PIKfyve-TRPML1-Myosin II axis regulates the speed of amoeboid cell migration

**DOI:** 10.1101/2024.01.22.575998

**Authors:** Philippe Dehio, Céline Michard, Juan Carlos Yam-Puc, Adrià-Arnau Martí i Líndez, Lucien Fabre, Thorsten Schaefer, Matthias P. Wymann, Klaus Okkenhaug, Thierry Soldati, Matthias Mehling, Christoph Hess

**Author notes:** Correspondence: Christoph Hess.

## Abstract

Amoeboid cell migration is key to efficient T cell immunity. Spatial polarization of organelles within cells, including endo-lysosomes, is a prerequisite of migration. However, how ultrastructural polarization is linked to the signaling requirements governing T cell migration, remains unknown. Here we show that signaling molecules generated by endo-lysosome-localized kinases regulate velocity of amoeboid migration. Specifically, imaging of T cells identified accumulation of endo-lysosomes decorated with the lipid kinases VPS34–PIKfyve at the uropod of polarized cells. Activity of VPS34 and PIKfyve regulated speed, but not directedness, of migrating T cells. Mechanistically, PI(3,5)P_2_ generated by the sequential action of VPS34 and PIKfyve mediated Ca^2+^ efflux from lysosomes via the mucolipin TRP cation channel 1 (TRPML1), thus controlling activity of myosin IIA and hence the generation of propulsive force through retrograde actin flow. The VPS34–PIKfyve kinases also regulated velocity of myeloid cells, as well as of the amoeba *Dictyostelium discoideum* – establishing the axis as an evolutionary conserved *speed control system* of amoeboid cell migration.

**Graphical Abstract:**

- The VPS34–PIKfyve axis is active on endo-lysosomes at the uropod of migrating T cells.
- VPS34 and PIKfyve promote myosin IIA activation and retrograde action flow.
- Amoeboid cell migration speed is controlled by VPS34 and PIKfyve via TRPML1.
- Regulation of amoeboid migration speed is a conserved function of the VPS34–PIKfyve axis.

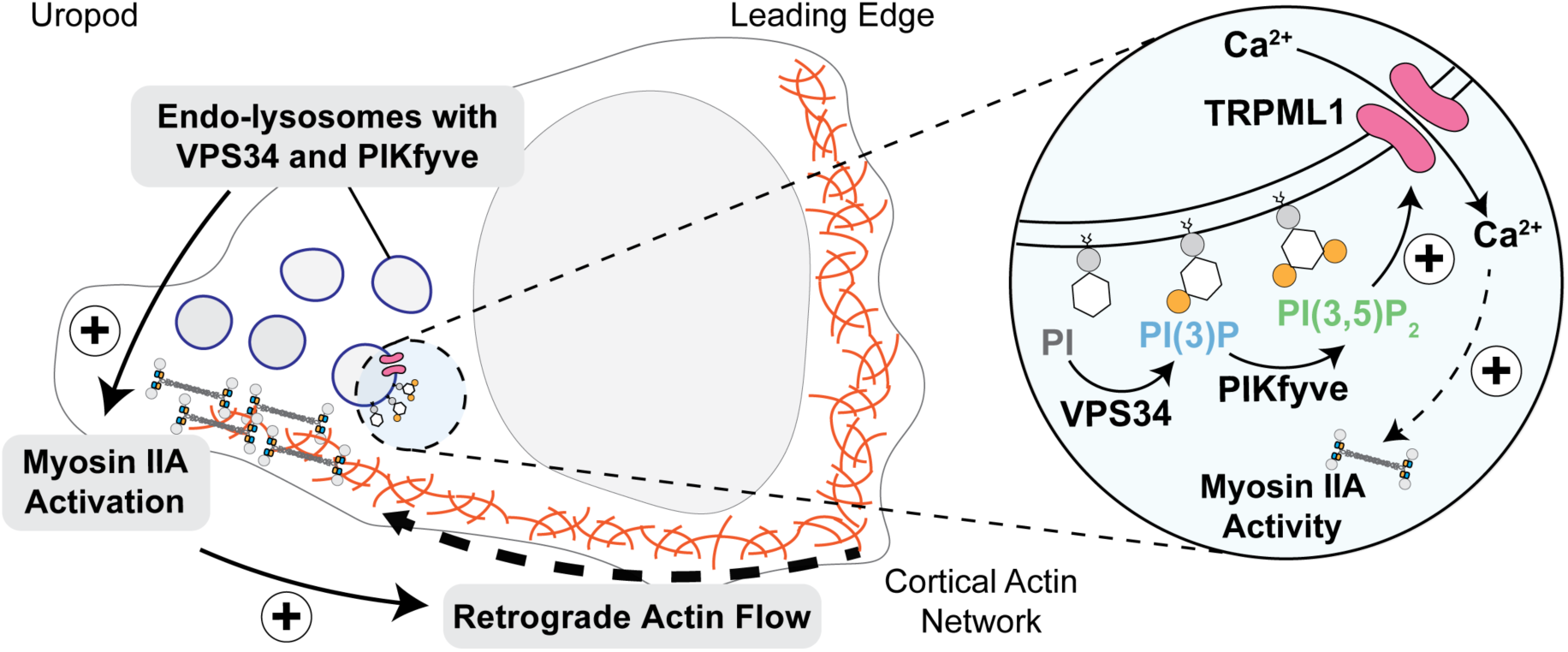

## Introduction

Cell migration is critically involved not only in immunity, but also early embryonic development, tissue homeostasis and regeneration. Across the eukaryotic domain, crawling-like, or amoeboid, movement is the most prevalent mode of cellular motility, with origins predating the emergence of metazoans.^1^ Amoeboid locomotion is characterized by bleb- or pseudopod-based membrane protrusion at the leading edge and retraction of the uropod – thereby establishing the direction of motion.^2–5^ At the molecular level, actomyosin contractility at the uropod, and actin polymerization at the leading edge, drive a retrograde actin flow, which propels cells forward when transmitted to the extracellular space.^6–10^ This directed motion of F-actin is regulated via phosphorylation of the non-muscle myosin IIA regulatory light chain (RLC).^11^ RLC is phosphorylated, and hence activated, by myosin light chain kinase (MLCK) in a calmodulin and cytosolic calcium (Ca^2+^) dependent process.^12,13^ Generation of propulsive force in amoeboid migration is accompanied by, and relies on, polarization of the cellular ultrastructure. In T cells, for instance, mitochondria translocated to the uropod have been shown to locally provide the necessary energy for myosin IIA activation.^14,15^ In dendritic cells (DCs), endo-lysosomes positioned at the uropod were demonstrated to support activation of myosin IIA by facilitating lysosomal Ca^2+^ release through the Ca^2+^-channel TRPML1.^16,17^ Binding of chemokines to their respective receptors triggers engagement of Rac1/2 and RhoA GTPase activity, which is affecting cell polarity and actin polymerization.^18,19^ The downstream signaling events that regulate amoeboid cell migration remain unexplored.

Here we investigated the role of lysosomes as signaling hubs in controlling amoeboid migration of T cells. Broader relevance of key findings was explored in non-related cell systems, including the non-metazoan organism *Dictyostelium discoideum* (*D. discoideum*).

## Results

### Endo-lysosomes re-locate together with VPS34-PIKfyve to the uropod of migrating T cells

To assess the localization of organelles among resting *vs.* chemokine-activated naïve murine CD8^+^ T cells, focused ion beam scanning electron microscopy (FIB-SEM) imaging was performed. Non-stimulated T cells were roughly circular in shape, and organelles distributed around the nucleus. Activation with the chemokine CCL19 induced cells to elongate and organelles to group towards one side of the cell (=polarization), including those with morphologic characteristics of endo-lysosomes (**Fig. 1A**). Immunofluorescence imaging of endo-lysosomes, using antibodies targeting Rab7 (late endosome and lysosome marker) and Lamp1 (lysosome marker), confirmed endo-lysosomal polarization towards one side of CCL19 activated naïve murine CD8^+^ T cells (**Fig. 1B**, top panels and lower left panel). Stimulation with CCL19 furthermore increased colocalization of Rab7 and Lamp1, indicative of endo-lysosomal maturation (**Fig. 1B**, lower right panel). Live-cell imaging of migrating naïve murine CD8^+^ T cells stained with a lysosomal dye also captured polarization of (endo)-lysosomes within cells, and established that endo-lysosomes accumulated at the uropod (**Fig. S1A**). Lastly, imaging naïve murine CD8^+^ T cells migrating in fluorescently tagged CCL19 (DY-649P1-CCL19) revealed focal accumulation of labelled CCL19 in intracellular vesicular structures that maintained their position throughout an observation period of up to five minutes (**Fig. 1C**). Immunofluorescence imaging confirmed accumulation of CCL19 in Rab7^+^ Lamp1^+^ positive regions, i.e. endo-lysosomes, in polarized naïve murine CD8^+^ T cells (**Fig. S1B**).

**Figure 1:**
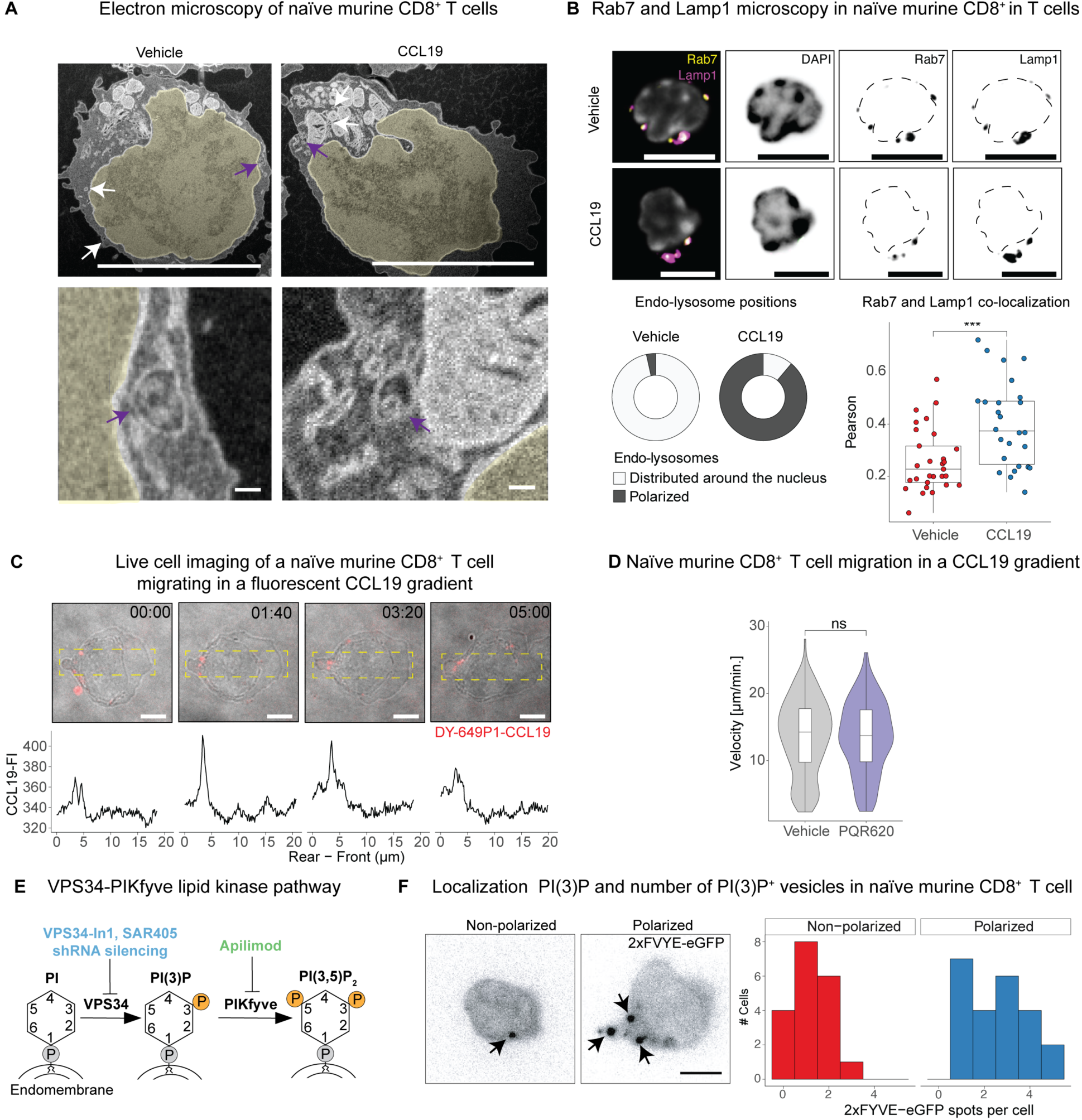
Endo-lysosomes decorated with VPS34–PIKfyve relocate to the uropod of polarized T cells |. **A**) FIB-SEM microscopy images of naïve murine CD8^+^ T cells treated with vehicle control (*left panels*) or 0.5 µg / ml CCL19 (*right panels*). Arrows indicate endo-lysosomes, further magnified examples are pointed out in purple (lower panels). The nucleus is pseudo-colored in yellow. *Scale bar: 5 µm for top panels, 100 nm for lower panels.* **B**) Immunofluorescence imaging of endo-lysosomes in naïve murine CD8^+^ T cells treated with either vehicle control or 0.5 µg / ml CCL19. *Top panels*: Representative images of a cell stained with DAPI, Rab7, and Lamp1. The dashed line in Rab7 and Lamp1 panels indicates the outline of the nucleus. *Bottom left panel:* Fraction of cells with polarized or non-polarized endo-lysosomes assessed via Rab7 staining in presence (n = 45) or absence (n = 29) of CCL19. *Bottom right panel:* Colocalization of Rab7 and Lamp1 fluorescence signal in naïve murine CD8^+^ T cells after treatment with vehicle control (n = 29) or CCL19 (n = 26). *Scale bar: 5 µm.* **C**) *Top panels*: Bright field images merged with DY-649P1-CCL19 signal (red) of a representative naïve murine CD8^+^ T cell. Lower panels: Quantification of the CCL19 signal fluorescent intensity (FI) from the above image along the x-axis of the dashed box. *Scale bar: 5 µm.* **D**) Velocity of naïve murine CD8^+^ T cells in a CCL19 gradient, following treatment with vehicle (n = 272) or the mTOR inhibitor PQR602 (n = 232). **E**) Diagram of the VPS34-PIKfyve pathway with inhibitors used (in color) and their respective targets. VPS34 and PIKfyve phosphorylate the inositol ring of phosphatidylinositol (PI) at the third and fifth position, respectively (represented by an orange circle marked with a “P”). **F**) Representative images of naïve murine CD8^+^ T cells transfected with the PI(3)P-sensor 2xFYVE-eGFP, showing quantification of detected PI(3)P^+^ vesicles (= 2xFYVE-eGFP spots) in polarized and non-polarized migrating naïve CD8^+^ T cells (n = 42). Data are pooled from three independent experiments. Arrows indicate PI(3)P^+^ vesicles. *Scale bar: 5 µm. Panels B-D and F: Representative data from three or more independent experiments. Not significant = p-value (p) > 0.05, *** p < 0.001; Wilcoxon rank-sum test for panels B and D*.

Lysosomes function as signaling hubs for various kinases, including the mechanistic target of rapamycin (mTOR). Specifically, mTOR, in conjunction with the Ragulator–Rag complex, integrates environmental signals (e.g. nutrients and cytokine signaling) on lysosomes.^20^ mTOR has been suggested to be activated by chemokine signaling, and regulate T cell migration.^21–23^ We thus reasoned that endo-lysosomes might control T cell migration by recruiting mTOR signaling to the uropod. Using the selective mTOR1/2 inhibitor PQR620, we blocked mTOR activity in naïve CD8^+^ T cells migrating in a CCL19 gradient.^24^ The mTOR blocking activity of PQR620 was confirmed by assessing phosphorylation of the mTOR target S6K by flow cytometry in activated CD8^+^ T cells (**Fig. S1C**). Inhibiting mTOR did not affect velocity of naïve murine CD8^+^ T cells in our system (**Fig. 1D**), possibly due to low mTOR activity in naïve T cells.^21–23,25^

However, endo-lysosomal membranes are also the target of the lipid-kinases VPS34 and PIKfyve, which phosphorylate phosphatidylinositol (PI) to PI(3)P and PI(3,5)P_2_, respectively (**Fig. 1E**).^26^ These kinases impact endo-lysosomal maturation, receptor recycling and Rac1 activity, features that have been associated with regulation of cell motility.^26–28^ This prompted us to consider their role in T cell migration. A fluorescent probe consisting of two PI(3)P-binding FYVE domains fused to an eGFP (2xFYVE-eGFP) enables subcellular localization of PI(3)P, i.e. the substrate of PIKfyve by fluorescence imaging.^29^ 2xFYVE-eGFP accumulated on vesicular structures at the uropod of CCL19-polarized naïve murine CD8^+^ T cells (**Fig. 1F**), thus capturing VPS34 activity in these cells and opening the possibility for involvement of the VPS34–PIKfyve axis in regulating amoeboid cell migration. This notion was further supported by a recently published CRISPR screen that identified *PIK3C3*, the gene encoding VPS34, as a regulator of T cell migration into the central nervous system in an experimental autoimmune encephalitis model.^30^

In all, these imaging studies (i) demonstrated that chemokine exposure of naïve CD8^+^ T cells induced endo-lysosomal maturation and polarization to the uropod, and (ii) provided evidence for engagement of lipid kinase activity on uropod-localized endo-lysosomes.

### T cell velocity, but not directedness, is controlled by VPS34 and PIKfyve

Using inhibitors of VPS34 and PIKfyve, we first explored how activity of these kinases related to accumulation of endo-lysosomes in polarized T cells. Inhibition of VPS34, using the potent and selective inhibitor VPS34-In1, was verified by imaging studies (**Fig. S2A**).^31^ Apilimod is a highly selective PIKfyve inhibitor in clinical development.^32,33^ Importantly, both inhibitors do not affect other lipid kinases, such as class 1 PI3K (e.g. PI3Kγ), which have been implicated in T cell migration.^34^ At 2 µM, i.e. the concentration used throughout, neither VPS34In-1 nor Apilimod affected the viability of murine CD8^+^ T cell blasts after 6 hours of treatment (**Fig. S2B**). With these tools in hand, we blocked VPS34 or PIKfyve in CCL19 stimulated naïve murine CD8^+^ T cells and examined cells using brightfield imaging. Inhibiting either kinase resulted in the appearance of large vesicles at the uropod of polarized cells (**Fig. 2A**, upper panel). The fluorescent signal from a lysosomal dye was also increased in VPS34In-1 as well as Apilimod-treated naïve murine CD8^+^ T cells, indicative of an increase in endo-lysosomal volume (**Fig. 2A**, lower panel). These findings align with the known role of the VPS34–PIKfyve system in endo-lysosomal homeostasis.^26^

**Figure 2:**
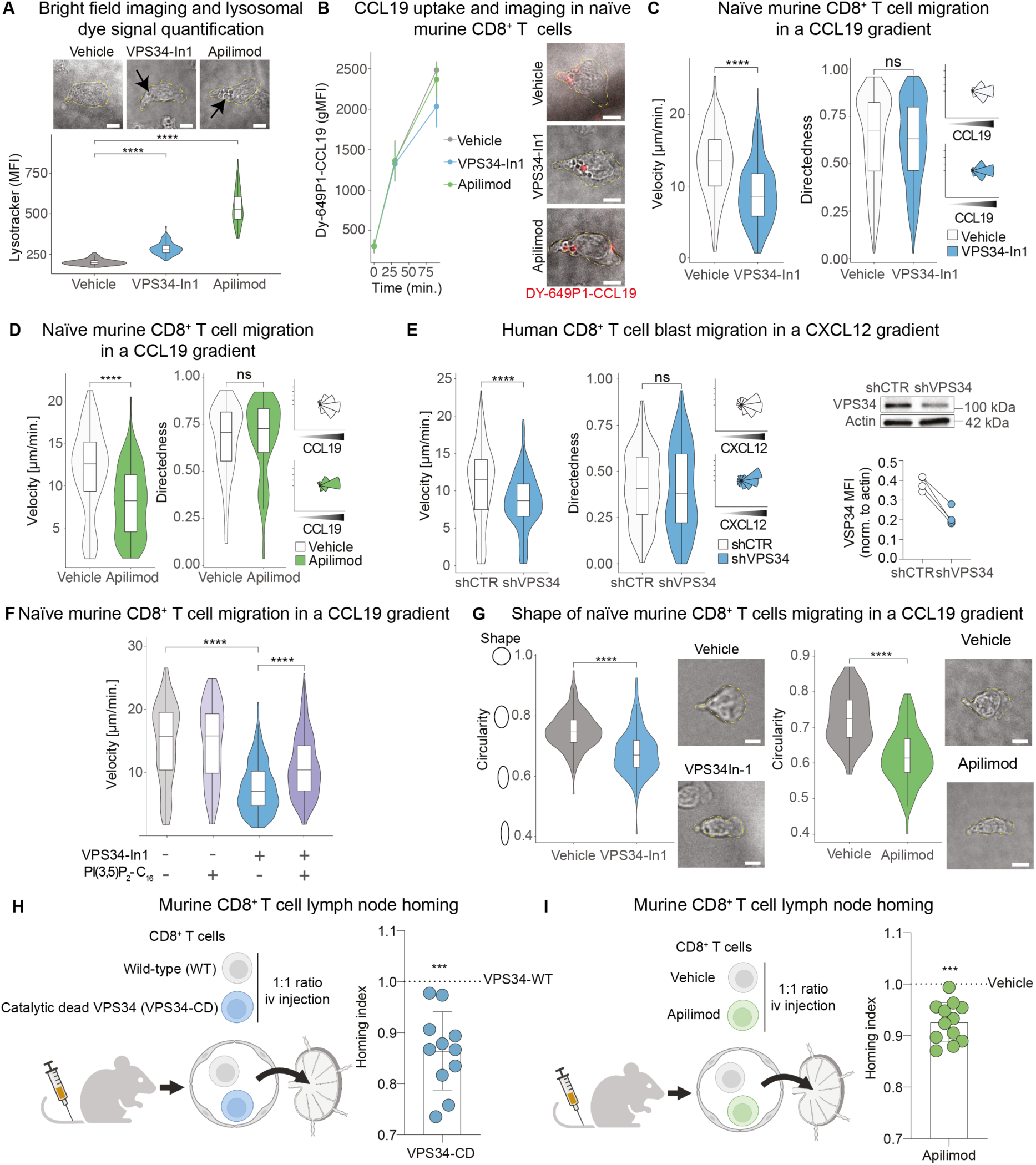
T cell velocity, but not directedness, is controlled by VPS34 and PIKfyve |. **A**) *Upper panel:* Representative bright field images of naïve murine CD8^+^ T cells migrating in a CCL19 gradient, treated with either vehicle control, VPS34-In1, or Apilimod. Arrows point at (large) vesicles. *Lower panel:* Quantification of lysosomal dye signal (mean fluorescent intensity, MFI) per cell, treated with vehicle control (n = 60), VPS34-In1 (n = 106), or Apilimod (n = 73). **B**) *Left panel*: Time course analysis of DY-649P1-CCL19 uptake in naïve murine CD8^+^ T cells (flow cytometry, geometric mean intensity (gMFI) and standard deviation of three biological replicates). *Right panel:* Representative bright field images merged with DY-649P1-CCL19 signal (red) of naïve murine CD8^+^ T cells migrating in a CCL19 gradient, treated with vehicle control, VPS34-In1, or Apilimod. **C**) Migration metrics (velocity, directedness, and angle of migration) of naïve murine CD8^+^ T cells treated with vehicle control (n = 362) and VPS34-In1 (n = 193), or **D**) vehicle control (n = 125), and Apilimod (n = 121). **E**) *Left panels:* Migration metrics of human CD8^+^ T cell blasts transduced with scrambled control (shCTR; n = 292), or VPS34-targeting shRNA (shVPS34, n = 255), and placed in a CXCL12 gradient. *Right panel:* VPS34 knockdown efficiency as determined by Western blotting; connections between dots link samples from the same biologic replicate. **F**) Velocity of naïve murine CD8^+^ T cells placed in a CCL19 gradient and treated with vehicle control (n = 175), PI(3,5)P_2_-C_16_ (n = 149), VPS34-In1 (n = 466), or PI(3,5)P_2_-C_16_ & VPS34-In1 (n = 636). **G**) Circularity analysis and representative bright field images of naïve murine CD8^+^ T cells migrating in a CCL19 gradient. *Left panel:* Vehicle control (n = 161) and VPS34-In1 (n = 157). *Right panel:* Vehicle control (n = 40) and Apilimod (n = 102). **H**) Schematic of experiment and result of CD8^+^ T cell homing analysis, comparing transfer of wild-type vs. catalytically-inactive VPS34 (pooled data from two independent experiments with 5-6 recipient animals each). **I**) Comparison of vehicle vs. Apilimod-treated CD8^+^ T cells in homing experiments (pooled data from two independent experiments with 5 recipient animals each). *Yellow dashed lines indicate cell outline, and the scale bars indicate 5 µm in panels A, B, and G. Panels A - G: Representative data from three or more independent experiments. ns = p > 0.05, *** p < 0.001, **** p < 0.0001; Wilcoxon rank-sum test for panels A, C-E, G-I; Aligned Rank Transform (ART) ANOVA for panel F*.

To test whether uptake and vesicular accumulation of chemokines at the uropod was affected by inhibition of VPS34 or PIKfyve, naïve murine CD8^+^ T cells were activated using fluorescent CCL19 in presence or absence of VPS34In-1 or Apilimod, respectively. Neither inhibition with VPS34In-1 nor Apilimod affected uptake of CCL19, as measured by (i) flow cytometry after 30 or 90 minutes (**Fig. 2B**, left panel), and (ii) accumulation of fluorescence in vesicular structures at the uropod (**Fig. 2B**, right panel). This suggested that the uptake of chemokine and cell polarization were not affected by inhibition of the VPS34–PIKfyve axis. We then went on and tested how inhibiting VPS34 or PIKfyve impacted T cell motility, using an under-agarose migration assay (**Fig. S2C**). Inhibition of VPS34 and PIKfyve both reduced the migration velocity of naïve murine CD8^+^ T cells, while not altering directedness and angle of migration towards the source of chemokine (**Fig. 2C**,**D**, **Fig. S2D**). VPS34In-1 and Apilimod also reduced the speed of murine CD8^+^ T cell blasts in a CCL19 gradient (**Fig. S2E**). shRNA-mediated silencing of VPS34 in human CD8^+^ T cell blasts likewise decreased their speed, while preserving directedness and angles of migration (**Fig. 2E**). Finally, the structurally non-related, selective VPS34 inhibitor SAR405 phenocopied migration deficits induced by VPS34 and PIKfyve inhibition in human CD8^+^ T cell blasts migrating in a CXCL12 gradient (**Fig. S2F**).^35^

Assessing whether the product of the VPS34–PIKfyve axis was indeed responsible for the observed phenotype, cell culture medium was supplemented with dipalmitoyl-PI(3,5)P_2_. Evidencing the importance of PI(3,5)P_2_, reduced migration velocity imposed by inhibition of VPS34 in naïve murine CD8^+^ T cells was partially rescued by addition of this synthetic PI(3,5)P_2_ (**Fig. 2F**). Rapidly migrating T cells tend to be elongated and less cirucluar.^6^ Notably, inhibition of VPS34 and PIKfyve, and thus slowing their migration, also decreased the average circularity of naïve murine CD8^+^ T cell migrating in a CCL19 gradient (**Fig. 2G**). This phenotypic constellation (decreased migration speed among less circular cells) was compatible with insufficient myosin IIA activity in VPS34–PIKfyve inhibited cells – a notion further explored below.^4,36,37^ To test whether also *in vivo* migration of CD8^+^ T cells was impacted by VPS34 and PIKfyve, adoptive transfer experiments were performed. Specifically, murine CD8^+^ T cells expressing either wild-type or a catalytic inactive form of VPS34 were injected in a 1:1 ratio into the tail vein of recipient mice, and their homing into lymph nodes (inguinal, axillary, cervical) was quantified (**Fig. 2H**, left panel).^38^ In these experiments, the number of wild-type CD8^+^ T cells migrating into lymph nodes was consistently higher than that of cells expressing catalytic inactive VPS34 (**Fig. 2H**, right panel). Likewise, pre-treatment of murine CD8^+^ T cells with Apilimod reduced lymph node homing relative to vehicle-treated control cells (**Fig. 2I**). Frequencies of transferred CD8^+^ T cells in the spleen were similar to input, irrespective of VPS34 and PIKfyve sufficiency *vs.* insufficiency (**Fig. S2G**,**H**) – aligning with passive accumulation in the red pulp and marginal zone compartment.^39,40^

Together these experiments identified the lipid kinase pathway PI→PI(3)P→PI(3,5)P_2_, governed by VPS34 and PIKfyve, as a modifier of endo-lysosomal turnover at the uropod of T cells, while leaving chemokine uptake unaltered. More importantly, activity of these kinases, through their end product, determined velocity of migrating CD8^+^ T cells, without affecting chemotactic directedness.

### VPS34 and PIKfyve regulate retrograde actin flow and myosin IIA activation

The flow of actin – i.e. the propulsion force underlying amoeboid cell migration – relies on synchronized actin polymerization at the leading edge, and myosin IIA-induced F-actin traction at the uropod.^6,7,9,10^ The importance of myosin IIA activity in governing cell migration was specifically verified for T cells by using two distinct approaches. First, direct (blebbistatin), as well as indirect (ML-7), inhibition of myosin IIA activity reduced migration speed of human CD8^+^ T cell blasts as well as human naïve CD8^+^ T cells, irrespective of upstream VPS34 activity (**Fig. 3A**-**C**). Second, nocodazole, which increases myosin IIA activity, increased velocity of migration both in presence and absence of VPS34 blockade, and also restored decreased circularity induced by inhibition of VPS34 to baseline levels (**Fig. 3A**, **Fig. 3D** and **Fig. S3A**).^41–45^ Given these clear data, we hypothesized that the VPS34–PIKfyve system impacted migration via regulating the activity of myosin IIA – an alternative possibility being an impact of the kinase-system on actin polymerization. The latter idea was refuted by the finding that actin polymerization in CCL19-treated naïve murine CD8^+^ T cells was unaffected upon inhibition of either VPS34 or PIKfyve (**Fig. 3E**). By contrast, in this same system myosin IIA activity – read out by the phosphorylation status of RLC – was diminished upon blocking of either kinase (**Fig. 3F**), also aligning with elongation observed among T cells migrating in presence of VPS34 or PIKfyve inhibitors (**Fig. 2G**).

**Figure 3:**
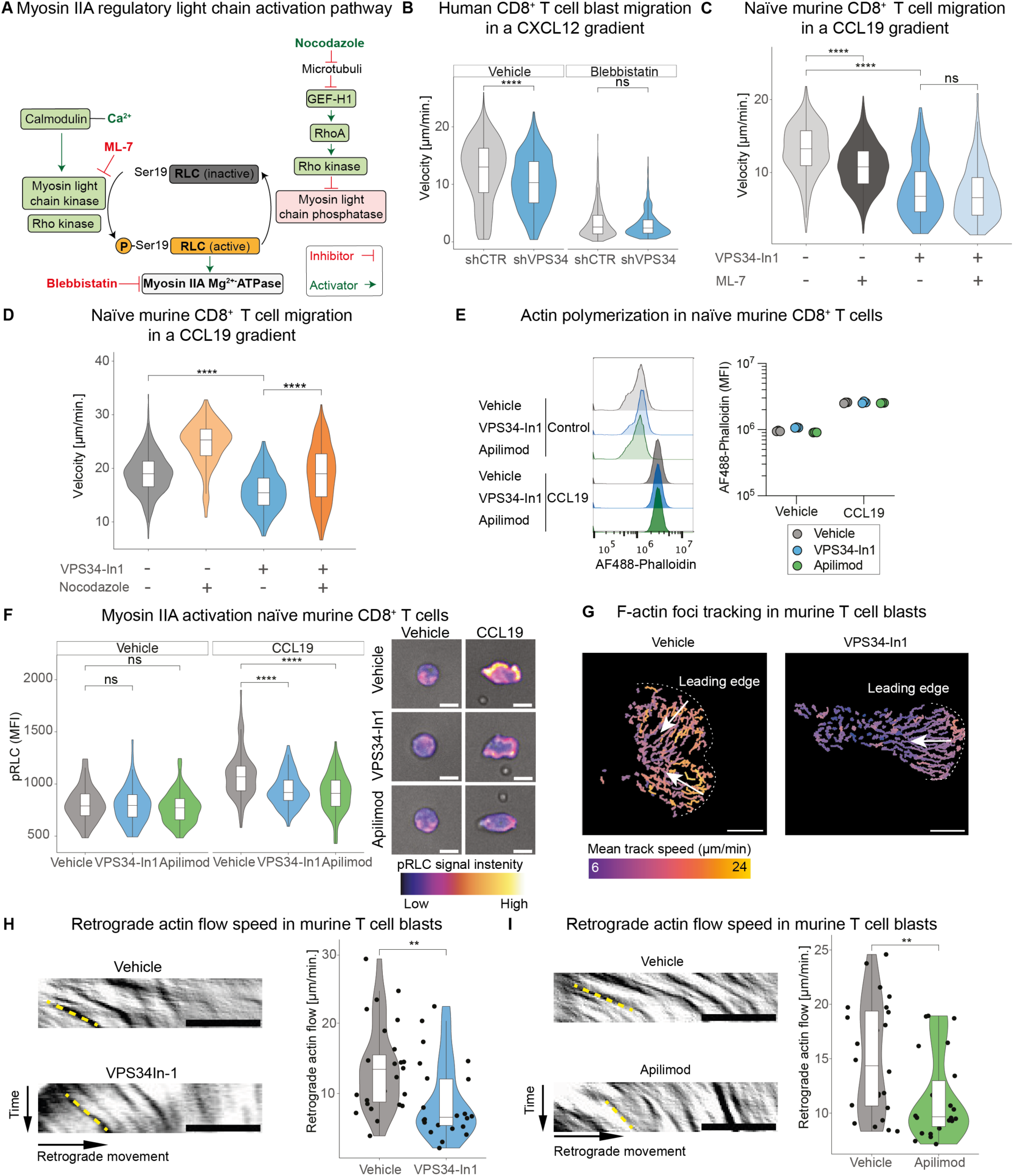
VPS34 and PIKfyve regulate retrograde actin flow and myosin IIA activation|. **A**) Schematic of the myosin IIA regulatory light chain activation pathway. Inhibitors or inhibitory interactions are shown in red, activators or activating interactions in green. **B**) Velocity of human CD8^+^ T cell blasts transduced with control shRNA (n = 1037 for vehicle control; n = 759 for Blebbistatin) or VPS34-shRNA (n = 534 for vehicle control; n = 594 for Blebbistatin), placed in a CXCL12 gradient in presence vs. absence of Blebbistatin. **C**) Velocity of naïve murine CD8^+^ T cells in a CCL19 gradient treated with vehicle control (n = 815), ML-7 (n = 1019), VPS34-In1 (n = 199), and ML-7 & VPS34-In1 (n = 238); or **D**) Vehicle control (n = 439), Nocodazole (n = 185), VPS34-In1 (n = 243), and Nocodazole & VPS34-In1 (n = 336). **E**) *Left panel*: Representative flow cytometry histograms of F-actin staining (AF488-Phalloidin) in naïve murine CD8^+^ T cells assessed one minute after treatment with vehicle control or CCL19. *Right panel:* Phalloidin mean fluorescence intensity (MFI, flow cytometry) of three technical replicates. **F**) *Left panel:* Per cell MFI signal (fluorescence microscopy) among naïve murine CD8^+^ T cells stained for myosin IIA RLC phosphorylated at serine 19 (pRLC) and treated with vehicle control (no CCL19: n = 81; CCL19: n = 75), VPS34-In1 (no CCL19: n = 79; CCL19: n = 88), and Apilimod (no CCL19: n = 139; CCL19: n = 64). *Right panels:* Representative bright-field images of cells overlaid with pseudo-colored pRLC signal. **G**) Representative tracks of F-actin foci derived from actin-reporter T cell blasts with or without VPS34-In1, pseudo-colored according to speed. Dashed lines denote the leading edge, arrows indicate the direction of actin flow. **H**) Kymograph-based speed analysis of retrograde actin flow (*right panels*) in actin-reporter T cell blasts treated with vehicle control (n = 27) or VPS34-In1 (n = 24), and **I**) vehicle control (n = 25) or Apilimod (n = 26). *Left panels*: Representative kymographs of F-actin reporter signal along the cell’s axis over a 30 sec period. The yellow dashed line indicates an exemplary actin trace used for speed analysis. *Scale bars in panels F-I represent 5 µm*. *Panels B-I represent results from three independent experiments. ns = p > 0.05, ** p < 0.01, **** p < 0.0001; Wilcoxon rank-sum test for panels F, H, I; ART ANOVA for panels B - D*.

To dynamically visualize dependency of actin flow on the VPS34–PIKfyve system, T cell blasts from Lifeact-GFP expressing mice were confined by serum-free agarose on a non-adhesive coating and activated by CCL19 contained in the polymer (**Fig. S3B**).^46^ As expected, inhibition of either VPS34 or PIKfyve reduced the speed of cortical F-actin foci moving within cells (**Fig. 3G**-**I**). Collectively, these findings established that VPS34 and PIKfyve regulated retrograde actin flow – the main propulsion mechanism for T cell migration – via tuning activity of myosin IIA.

### VPS34 and PIKfyve regulate migration velocity of T cells via lysosomal Ca^2+^

We next aimed to explore how activity of the VPS34–PIKfyve axis, myosin IIA activity, and T cell migration were mechanistically interlinked. Endo-lysosomes function as Ca^2+^ stores, and cytosolic Ca^2+^ promotes activation of myosin IIA (**Fig. 3A**).^16,17^ We therefore imaged intracellular Ca^2+^ in naïve murine CD8^+^ T cells migrating in a CCL19 gradient in presence or absence of inhibitors for VPS34 and PIKfyve, respectively. Inhibition of either kinase resulted in a notable increase in lysosomal Ca^2+^ (**Fig. 4A**). Substantiating specificity of this signal, chelating intracellular Ca^2+^ with BATPA-AM caused progressive fading of the calcium indicator in lysosomes (**Fig. 4B** and **Fig. S4A**).

**Figure 4:**
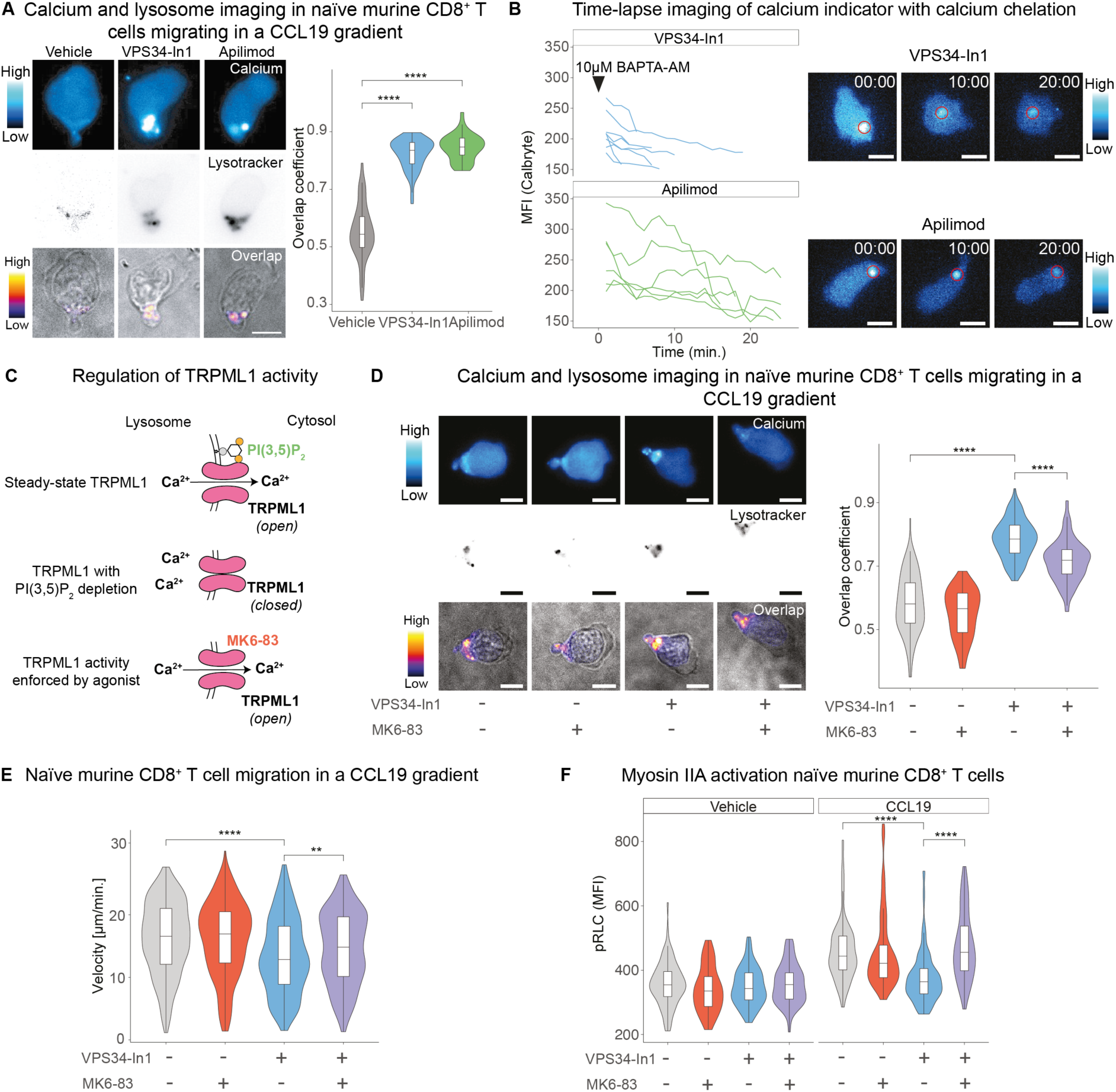
VPS34 and PIKfyve regulate migration velocity of T cells via lysosomal Ca^2+^ |. **A**) *Left panels:* Fluorescence images of naïve murine CD8^+^ T cells stained with a calcium indicator dye (Calbryte) and a lysosomal dye (LysoTracker). Bright field images were merged with a pseudo color map indicating the overlap of calcium and lysosomal signals (Pearson correlation coefficient). *Right panel:* Overlap coefficient (= colocalization) of calcium and lysosomes in cells treated with vehicle control (n = 45), VPS34-In1 (n = 44), or Apilimod (n = 37). **B**) *Left panel:* Time-course analysis of calcium indicator signal (MFI) in large vesicles induced by VPS34-In1 (n = 8) or Apilimod (n = 6) after BAPTA-AM addition in naïve murine CD8^+^ T cells. *Right panels:* Representative calcium indicator image of tracked vesicle*s,* marked by a red circle. **C**) Schematic of TRPML1 activation: Green symbols denote the TRPML1 agonist PI(3,5)P_2_ and MK6-83. **D**) *Left panels:* Fluorescence images of naïve murine CD8^+^ T cells stained with a calcium indicator dye (Calbryte) and a lysosomal dye (LysoTracker). Bright field images were merged with a pseudo color map indicating the overlap of calcium and lysosomal signals (Pearson correlation coefficient). *Right panel:* Overlap coefficient (= colocalization) of calcium and lysosomes in cells treated with vehicle (n = 94), VPS34-In1 (n = 120), MK6-83 (n = 65) or VPS34In-1 & MK6-83 (n = 122). **E**) Velocity of naïve murine CD8^+^ T cells migrating in a CCL19 gradient, treated with vehicle control (n = 560), VPS34-In1 (n = 408), MK6-83 (n = 418), and a combination of VPS34-In1 & MK6-83 (n = 580). **F**) Per cell MFI signal (fluorescence microscopy) among naïve murine CD8^+^ T cells stained for pRLC and treated with vehicle control (no CCL19: n = 83; CCL19: n = 110), VPS34-In1 (no CCL19: n = 72; CCL19: n = 113), MK6-83 (no CCL19: n = 47; CCL19: n = 72), and VPS34-In1 (no CCL19: n = 71; CCL19: n = 74). *Scale bars in panels A and B indicate 5 µm. Panels A, B, D-F are representative results of 3 independent experiments. ** p < 0.01, **** p < 0.0001; Wilcoxon rank-sum test for panel A; ART ANOVA for panel D-F*.

The flux of Ca^2+^ from lysosomes to the cytoplasm is gated by TRPML1, a process regulated by PI(3,5)P_2_ (**Fig. 4C**).^47–49^ We thus hypothesized that inhibition of VPS34 or PIKfyve, and thus depletion of PI(3,5)P_2_, hinders lysosomal Ca^2+^ efflux – leading to accumulation of lysosomal Ca^2+^ in migrating naïve murine CD8^+^ T cells. Functional engagement of TRPML1 should thus restore lysosomal Ca^2+^ efflux and reverse VPS34-dependent migration deficits in T cells. To test this idea we monitored migration of naïve murine CD8^+^ T cells in presence or absence of VPS34-In1 and/or the specific and potent TRPML1 agonist MK6-83 (**Fig. 4C**).^50^ Indeed, MK6-83 reduced lysosomal Ca^2+^ sequestration and partially rescued the speed deficit imposed by inhibition of VPS34 (**Fig. 4D**,**E**). TRPML1 activation also reverted reduced myosin IIA activation imposed by inhibition of VPS34 (**Fig. 4F**).

From these experiments a molecular framework emerged, where VPS34–PIKfyve derived PI(3,5)P_2_ was regulating TRPML1 mediated lysosomal Ca^2+^-efflux, thereby triggering the calmodulin–MLCK pathway and hence activation of myosin IIA.

### Regulation of amoeboid migration is a conserved function of the VPS34–PIKfyve axis

To explore whether the VPS34–PIKfyve system is involved in regulating amoeboid migration also in other cell types, HL-60-derived neutrophils and murine bone marrow-derived DCs were used. Both, neutrophils in the presence of fMLP and DCs exposed to CCL19 migrated slower when blocking either VPS34 or PIKfyve (**Fig. 5A** and **Fig. 5B**, left panel). Infiltration of DCs under an agarose-matrix was likewise reduced when blocking VPS34 or PIKfyve (**Fig. 5B**, right panel and **Fig. S5A**). These findings expanded the role of the VPS34–PIKfyve system in regulating velocity of amoeboid cell migration beyond T cells. Of note, VPS34 and PIKfyve orthologues are widely expressed across all eukaryotic kingdoms (**Fig. S5B**).^51^ We thus considered the idea that the VPS34–PIKfyve → TRPML1 pathway could be a conserved biological module regulating velocity of amoeboid cell migration. To test this idea, we used the social amoeba and professional phagocyte *D. discoideum*, a non-metazoan model organism that separated from the animals and fungi a billion years ago. *D. discoideum* exhibit similar vesicular alterations upon PIKfyve inhibition as metazoan cells, and its migration is myosin II-dependent.^52,53^ Further, absence of the *D. discoideum* TRPML1 orthologue Mucolipin (MCLN) impairs Ca^2+^-release from endo-lysosomes.^54^ Indeed, spontaneous amoeboid migration of PIKfyve knock-out (ko) and Apilimod-treated wild-type (Ax2) *D. discoideum* cells was slower than that of respective control cells. Inhibition of PIKfyve with Apilimod did not alter the speed of migration among PIKfyve ko cells, arguing for the specificity of the inhibitor (**Fig. 5C**). Moreover, MCLN-deficient *D. discoideum* phenocopied PIKfyve deficiency (**Fig. 5D**). Supporting the notion that MCLN was the target of PI(3,5)P_2_ also in *D. discoideum* cells, the alpha-fold model of MCLN displayed a positively charged pocket similar to the PI(3,5)P_2_ binding pocket in the N-terminus of TRPML1 (**Fig. 5E**).^47,48^ In line, 5 out of 7 amino acids in this poly-basic domain are conserved between TRPML1 and MCLN (**Fig. S5C**), and *in silico* docking analyses suggested potential binding of PI(3,5)P_2_ to this pocket in MCLN (**Fig. 5E**). To explore whether regulation of amoeboid migration speed has evolved as an early feature of the VPS34–PIKfyve axis, we extracted the number proteins with zinc finger FYVE domains from the public domain.^55^ PI(3)P binding to the FYVE zinc finger domain regulates function and recruitment of proteins containing this motif. Therefore, the number of distinct proteins with FYVE domains may serve as a proxy for a potential functional involvement of the VPS34–PIKfyve kinase system. While the number of proteins harboring this motif is relatively low in unicellular organisms (including *D. discoideum*), it increases in metazoans along with complexity (**Fig. S5D**). It hence is plausible that regulation of migration speed is an ancient function of the VP34–PIKfyve axis.

**Figure 5:**
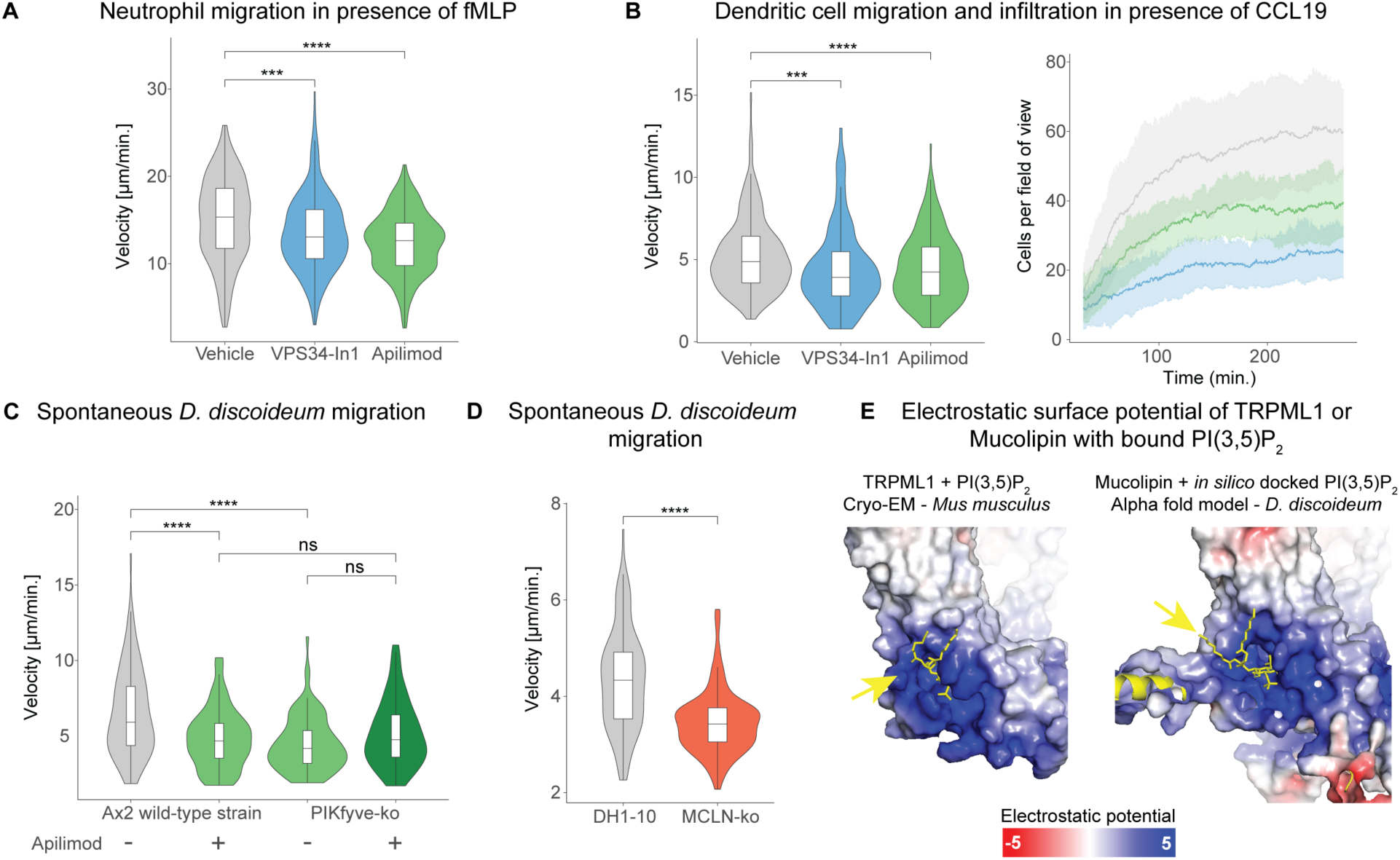
Regulation of amoeboid migration is a conserved function of the VPS34–PIKfyve pathway|. **A**) Velocity of HL-60 derived neutrophils treated with vehicle control (n = 214), VPS34-In1 (n = 243), and Apilimod (n = 194), migrating in presence of the chemokine fMLP. **B**) *Left panel*: Velocity of DCs migrating in CCL19, exposed to vehicle control (n = 412), VPS34-In1 (n = 104), Apilimod (n = 205). *Right panel*: Mean and standard deviation of number of cells detected per fields of view (n = 6 per condition) infiltrating under the agarose matrix. **C**) Velocity of *D. discoideum* wild-type (Ax2) cells treated with vehicle control (n = 124) or Apilimod (n = 87), and PIKfyve-orthologue ko cells treated with vehicle control (n = 91) or Apilimod (n = 74). **D**) Velocity of DH1-10 (wild-type strain, n = 113) and mucolipin-ko (MCLP-ko, n = 114) *D. discoideum* cells. **E**) *Left panel*: Experimentally determined structure of TRPML1 (public domain) with bound PI(3,5)P_2_.^47^ *Right panel*: Predicted structure of Mucolipin with PI(3,5)P_2_ docked *in silico*. The structures are aligned and superimposed with electrostatic potential. *Panels A-D: Representative results from three or more independent experiments. ns p > 0.05, *** p < 0.001, and **** p < 0.0001; Wilcoxon rank-sum test used in panels A, B and D; ART ANOVA for panel C*.

In all, these findings suggested an evolutionarily conserved mechanism in which PI(3,5)P_2_, generated by the VPS34–PIKfyve axis, regulated TRPML1 activity, thereby controlling the release of Ca^2+^ from lysosomes at the uropod of migrating cells – and hence velocity of amoeboid cell migration.

## Discussion

The key finding of this study was that the VPS34–PIKfyve axis, in association with endo-lysosomes, locates to the uropod of migrating T cells. This lipid kinase pathway is thereby ideally positioned to regulate retrograde actin flow by enabling Ca^2+^ efflux from endo-lysosomes through TRPML1, and hence myosin IIA activity. Regulation of cell migration-velocity via this molecular axis applies to various cell types, including *D. discoideum*, i.e. unicellular eukaryotic cells.

Precise positioning of lysosomes has previously been shown to be important for amoeboid migration, ensuring site-specific availability of Ca^2+^ (Refs. ^16,17^). We now show that the VPS34– PIKfyve axis links local cytosolic availability of Ca^2+^ (i.e. endo-lysosomes) and Ca^2+^ appropriation from these stores to engage myosin IIA activity. An increasing understanding of how ultrastructure and function are mechanistically interlinked highlights the importance to interrogate the molecular mechanisms orchestrating context-specific spatial ultrastructural dynamics.

Microtubules, and the microtubule organizing center at the uropod of cells, are critically involved in shaping the ultrastructure of amoeboid migrating cells. In T cells, for example, mitochondria are coupled to microtubules via Miro-1, which enables their re-location to the uropod upon chemokine signaling.^15^ In an analogous manner, endo-lysosomes connected to microtubules by Arl8B or RILP/Rab7, have been shown to be re-positioned within cells depending on nutrient availability.^56,57^ It will be interesting to define how chemokine signaling and microtubule-dependent positioning of organelles are interlinked, and characterize the adaptor proteins that are involved in this process. Given the precise and dynamic ultrastructural adaptations happening in cells upon external input, further defining structure–function relationships of ultrastructural components, as well as their interactions, will also be relevant. At the uropod, mitochondria, endo-lysosomes, and the ER reside in close vicinity. Intriguingly, membrane contact sites physically connecting these organelles jointly regulate Ca^2+^ dynamics and provide microdomains for local signaling.^58–61^ Whether such contact sites selectively form as an ultrastructural adaptation in migrating cells and, if so, their biologic roles, remains to be investigated.

In our experiments with T cells, chemokine-induced F-actin formation was unaffected by VPS34–PIKfyve. Notably, in neutrophils and cancer cells, PIKfyve has been shown to stimulate activity of the GTPases Rac1/2, which control actin polymerization.^18,27,62^ It will be interesting to explore whether VPS34 and PIKfyve also control Rac1/2 in T cells and, if so, what the molecular and functional consequences of Rac1/2 activity are.

VPS34 and PIKfyve orthologues are universally expressed in eukaryotes and their appearance is known to predate the emergence of amoeboid cell migration.^51,63^ Roles previously assigned to the VPS34–PIKfyve axis are regulation of endocytic trafficking and of autophagy.^64^ The herein described role in governing the speed of amoeboid cell migration not only in vertebrate immune cells but also *D. discoideum*, indicates early functional assimilation of the VPS34–PIKfyve kinase system to also support this cellular process. Across taxa, F-actin polymerization and cortical actin-network contraction by myosin IIA lie at the center of amoeboid locomotion.^65,66^ This study thus implicates regulation of myosin IIA activation by lysosomal Ca^2+^ via VPS34–PIKfyve–TRPML1 axis as an evolutionary conserved regulator of this core mechanism.

In all, our work identified a conserved regulatory node linking local availability of Ca^2+^ stores (endo-lysosomes) and generation of lipid-mediators with the efflux of Ca^2+^ from their storage organelles. This spatially organized biologic system governs the speed of amoeboid cell migration.

## Acknowledgements

We thank the microscopy core facility of the Department of Biomedicine at the University and University Hospital of Basel for their technical support. CH was supported by the Swiss National Science Foundation (SNSF) (310030B_201277; 310030_192677; FZEB-0-180487) and the Novartis Foundation for Medical-Biological Research (NFMBR) (#23A070). PD was supported by a scholarship from the Swiss Academy for Medical Sciences (SAMW) and SNSF (183980), the NFMBR (#23A070), AlumniMedizin Basel, and the Freiwillige Akademische Gesellschaft Basel. We thank Daniel Legler for providing reagents.

## Author contributions

Conceptualization, C.H. and P.D.; Investigation, P.D., C.M., J.C.Y., A.M.L., and L.F.; Methodology, C.H., M.M., T.So., K.O., and P.D.; Resources C.H., M.M., T.Sc., M.P.W., K.O., and T.So.; Writing – Original Draft C.H. and P.D.; Writing – Review & Editing T.Sc., M.P.W., M.M., C.M. and T.So.; Supervision, C.H.; Project Administration C.H.

## Materials and Methods

### Cell isolation and culture

Cells and cell lines were cultured in RPMI1640 (containing 25 mM HEPES and L-Glut; Gibco, Cat. # 52400-025) + 10 % heat-inactivated fetal calf serum (hiFCS; Gibco, Cat. # 10270) + GlutaMax (diluted 100x; Gibco, Cat. # 35050-038) + 50 µM β-Mercaptoethanol (Gibco, Cat. # 31350-010) + 100 U/ml penicillin and streptomycin (Gibco, Cat. # 15140-122), referred to throughout the text as *complete medium*.

Naïve murine CD8^+^ T cells were isolated from the spleens and lymph nodes (inguinal, axillary, and cervical) from C57BL/6NCrl with a negative magnetic isolation kit, according to the manufacturer’s instruction (Stemcell, Cat. # 19858; Miltenyi Biotec, Cat. # 130-104-075). Mice of the age 6-25 weeks of both genders were used for the experiments. They were bred and housed in a specific-pathogen-free facility at the University of Basel. All experiments adhered to the guidelines of the Swiss Federal Veterinary Office and were approved by local authorities. Isolated murine T cells were cultured in 10 ng / ml recombinant murine IL-7 (Peprotech, Cat # 217-17) in complete medium and used up to 5 days after isolation.

HL-60 cells (a gift from Jürg Schwaller, University of Basel) were maintained in complete medium at a density of 2 x 10^5^ - 1.5 x 10^6^ cells / ml and split every 2-3 days. Neutrophil differentiation was induced by adding 1.3% (v/v) DMSO into complete medium. Differentiated cells were used 5-7 days after DMSO addition.

For DC differentiation, bone marrow was harvested from femurs and tibias of C57BL/6NCrl mice, and 2 x 10^6^ cells were plated in 10 cm Petri-dishes with 10 ml of complete medium containing 10 ng / ml GM-CSF (BioLegend, Cat. # 576306). At day 3 and 6 fresh medium with GM-CSF was added. On day 8, the floating fraction of cells (immature DCs) was harvested and frozen in 90 % hiFCS + 10 % DMSO and stored in liquid nitrogen. Frozen cells were thawed in a water bath at 37 ° C, added to prewarmed complete medium, and centrifuged at 200 x g for 5 min. The thawed cells were resuspended in complete medium & 10 ng / ml GM-CSF and 100 ng / ml LPS (Lucerna Chem, Cat. # abx082480) and matured for 14 h for migration experiments.

### Human CD8^+^ T cell isolation and shRNA-mediated VPS34 silencing

Peripheral Blood Mononuclear Cells (PBMCs) were isolated by density centrifugation from buffy coats of healthy and consenting donors (Blood donor center, University of Basel, male and female, 18-65 years old). CD8^+^ T cells were isolated using anti-CD8 microbeads, following the manufacturer’s instructions (Miltenyi Biotec, Cat. # 130-045-201). 2.5 x 10^6^ T cells were activated for two days with CD3/CD28-complexes (25 µl / ml, Stemcell, Cat. # 10971) and 100 U / ml of IL-2 (Bio-Techne Sales Corp., Cat. # 202-IL-050). The cells were then split into two 500 µl samples and treated with 100 µl of lentivirus solution. This solution contained 30 µl of scrambled or 10 µl VPS34-targeting shRNA per target sequence to a total of 30 µl and was supplemented with 6x polybrene to reach a final concentration of 6 µg / ml (Sigma Aldrich, Cat. # TR-1003-G) in a complete medium. The cells were spinoculated for 60 min at 1000 g at 30 °C. On day 3 after activation, the medium was replaced with complete medium containing 300 U/ml of IL-2. On day six post-activation, we sorted the cells for GFP^+^ expression using a FACSMelody Cell Sorter (BD). Sorted cells were expanded and maintained in complete medium containing 300 U/ml of IL-2 for up to two weeks before use in experiments. The high-titer shRNA lentivirus against human VPS34 (along with a scrambled control) was procured from VectorBuilder (shRNA 3 & 1 with EGFP; target sequences: GAGGCAAATATCCAGTTATAT, CCACGAGAGATCAGTTAAATA, GCTGGATAGATTGACATTTAG).

### Under-agarose cell migration assay

The under-agarose migration assay was adapted from a previously published method.^67^ Ibidi µ-Slide 8-well high Glass Bottom slides (Ibidi, Cat. # 80807) were coated with 150 µl of human plasma fibronectin (10 µg / ml; EMD Millipore, Cat. # FC010) in PBS and incubated at 37 °C for 2 h. The slides were then rinsed twice with 400 µl PBS per well. To cast the 1.2 % agarose gels, 0.48 g of ultra-pure agarose (Thermo Scientific, Cat. # 16500100) was mixed in 10 ml deionized water. Separately, 4 ml filtered hiFCS, 16 ml phenol-red free RPMI (Gibco, Cat # 11835-063), and 10 ml 2x HBSS (pH 7.2, from 10x HBSS stock; Gibco, Cat # 14065056) were combined and heated to 60 °C. The agarose was heated to a boil for three times in a microwave, interspersed with mixing. The medium, HBSS, and FCS mixture were then added to the molten agarose solution, followed by gentle mixing. Each well received 400 µl of the agarose mixture and was left to solidify with a closed lid. When inhibitors were used, 100 µl of a 5 x inhibitor solution in complete medium was added to the gel and incubated overnight at RT, sealed with parafilm. Before the migration assay, the medium solution was removed, and a well was created with a 3 mm biopsy puncher. Cells pretreated with inhibitors and stained with Hoechst 33342 (see below) were centrifuged at 200 x g for 5 min, and resuspended in complete, phenol-red free, inhibitor-containing medium at a concentration of 1 x 10^7^ cells / ml. 5.5 µl of the cell suspension was injected 2-3 mm away from the punched well using a micropipette with a gel-loading tip. All fluid was aspirated from the well, which was then filled with 20 µl of complete medium containing chemokines. For human CD8^+^ T cell blast migration assays, 2 µg / ml CXCL12 (Preprotech, Cat # 300-28A) was added, whereas for murine T cell migration assays, 2 µg / ml CCL19 (Peprotech, Cat # 250-27B) or 20 µM DY-649-P1-CCL19 (a kind gift from Daniel Legler, Biotechnology Institute Thurgau) in complete medium was added in the punched well.^68^ The dish was then transferred to the microscope. For DC experiments, the agarose gel was overlaid with 5 x inhibitor / vehicle solution and 5 x CCL19 solution (final concentration: 0.25 µg / ml) and cells were added to the well. For neutrophil migration assays, gels were overlaid with 5 x inhibitor / vehicle solution and 5x fMLP (Sigma-Aldrich, Cat # F3506, final concentration at 10 nM).

### Microscopy imaging modalities and analysis

Most imaging experiments (except *Dicty* cell migration assay) were conducted using Nikon inverted Ti/Ti2 microscopes (**Table 1**), equipped with a Photometrics 95B (25mm, back-illuminated sCMOS) camera. The Nikon Ti2 widefield microscope employed an LED, the Nikon Ti CSU-W1 spinning disk utilized a diode-pumped solid-state laser, and the Nikon Ti2 Cresty spinning disk used a Celesta solid-state laser as a light source. CFI Plan Apo Lambda objectives were used, with magnifications of 20x air (NA: 0.75), 40 x air (NA: 0.95), 100x air (NA: 1.45), and 60x water (NA: 1.2, Plan Apo VC). For live cell imaging, stage-top incubators maintained a constant temperature of 37 °C, 100 % humidity (20 l / h airflow), and 5 % CO_2_.

**Table 1.**

| Experiment | Microscope modality | System | Objective | Applied assay | Sampling rate |
| --- | --- | --- | --- | --- | --- |
| Immunofluorescence of endo-lysosomes | Widefield | Nikon Ti2 | 100x | Fixed and mounted cells |  |
| Immunofluorescence of RLC phosphorylation |  |  | 40x |  |  |
| Assessment of retrograde actin flow | Spinning disk confocal | Nikon Ti / Nikon Ti2 | 60x, 100x | Modified under-agarose (see below) | 1 fps |
| Live cell migration with inhibitors | Nikon Ti2 | Nikon Ti2 | 20x | Under-agarose | 20 or 30 sec |
| Live cell imaging: T cell morphology | Widefield | Nikon Ti2 | 40x & 1.5 optical zoom, 100x | Under-agarose |  |
| Live cell imaging: T cell with Lysosomal stains / | Widefield | Nikon Ti2 | 100x | Under-agarose | 10 fps or |
| DY-649P-CCL19 / 2x<br>FYVE-eGFP probe /<br>calcium indicator |  |  |  |  | single<br>images |

Cell morphology and fluorescence intensities were measured using FIJI. Regions of interest (ROI) were manually drawn in FIJI to obtain mean fluorescence intensities and shape descriptors. Each experiment involved analyzing at least three different fields of view per condition. Cell tracking, either for nuclei or brightfield images, was conducted using either Ilastik (with Pixel Classification followed by Tracking) or TrackMate in FIJI (Log detector; Simple LAP tracker with linking max distance: 20 microns, gap-closing max distance: 20 microns, gap-closing max frame gap: 2).^69,70^ Velocities and directedness were then calculated using a custom-written script in R. Colocalization analysis was performed on cropped images containing only one single cell using Huygens (Scientific Volume Imaging). For RAB7 and LAMP1 colocalization, a deconvolution step was performed in Huygens prior to the analysis.

### *Dictyostelium discoideum* culture and migration assay

The *D. discoideum* strains used in this study are listed in **Table 2**. Cells were axenically grown in 10 cm Petri dishes (Falcon) at 22°C in HL5 or HL5c medium (Formedium, Cat. # HLG0101) supplemented with 100 U/ml penicillin and streptomycin (Gibco, Cat. # 15140-122). At least 24 h before the migration experiments, all the cell lines were grown in HL5 medium. 6 x 10^3^ cells were seeded in each well of a 4-well µ-slide (Ibidi, Cat. # 80426) in HL5 medium and incubated 1 h at 22 °C. When necessary, 5 µM Apilimod (**Table 4**) or DMSO was added to the cells before the incubation. Time-lapse movies of 30 min with images taken at 30 sec intervals were recorded with a widefield inverted LEICA DMi8 microscope using a 40x objective. Images were processed with the Trackmate ImageJ plug-in.

**Table 2.**

| Genotype | Background strain | Source / Reference |
| --- | --- | --- |
| Wild-type | Ax2 | Buckley et al. 2019<sup>52</sup> |
| PIKfyve-ko |  |  |
| Wild-type | DH1-10 | Gift from Pierre Cosson; Lima et al. 2012<sup>54</sup> |
| Mucolipin-ko |  |  |

### Assessment of retrograde actin flow

The assessment of retrograde actin flow was carried out using a modified under-agarose assay (as described above). The slides were first plasma cleaned and PEG-coated at 4 °C overnight with 200µg/ml poly(L-lysine)-PEG (SuSoS, PLL(20)-g[3.5]-PEG(2)) in PBS. The slide was washed three times with PBS and 400 µL of 1.2 % agarose solution (without any serum). The gels were then treated overnight with 100 µL of 5x solutions of either the vehicle, VPS34In-1, or Apilimod in serum- and phenol-red free RPMI1640 containing 1.25 µg / ml CCL19. Subsequently, Lifeact T cell blasts were treated with either VPS34In-1 or Apilimod for 2 h and then injected under the agarose gel (as described above).

Actin flow videos were acquired at 1 frame per second (fps) for 30 s in the GFP channel (Ex.: 488nm) using an inverted spinning disk confocal (details above). The acquired images were then analyzed with FIJI: First, cells with visible actin spots were cropped and loaded into the KymographClear plugin with the average intensity projection.^71^ A line was then drawn through the cells, along the F-actin foci traces from the uropod to the leading edge, to generate a kymograph (with a line width of 5 pixels). The angle of a representative F-actin trace was analyzed, from which the speed was calculated by taking the tangent of the angle and scaling the result to µm / min. The tracking of F-actin foci was performed using Trackmate and the integrated LoG detector.

### Immunofluorescence staining

8-well removable cell culture dishes (ibidi, Cat. # 80841) were coated with 100 µg / ml Poly D-Lysine (Milipore Cat. # A-003-E) in PBS for 2 hours at 37 °C and washed three times with PBS. Then, 1 x 10^5^ naïve CD8^+^ T cells were plated per well, centrifuged for 30 s at 300 x g, and treated with 0.5 µg / ml CCL19 and inhibitors according to the protocol. At the end of the treatment, prewarmed EM-grade paraformaldehyde was added directly to the medium to achieve a final concentration of 4 %. The plate was then incubated for 15 minutes at 37 °C and washed once with PBS. Cells were permeabilized with 0.2% Triton X in PBS for 15 min and blocked with 5% goat serum in PBS-T (blocking buffer) for 1 hour. Primary antibodies were incubated overnight at 4 °C and secondary antibodies for 1 h in the dark at RT, with three intermittent washes with PBS-T. The antibodies were diluted in blocking buffer (**Table 3**). Finally, the slides were washed in PBS, mounted with a coverslip and ProLong Gold Antifade with DNA Stain DAPI (Invitrogen, Cat. # P36931) and sealed the next day with nail polish.

**Table 3.**
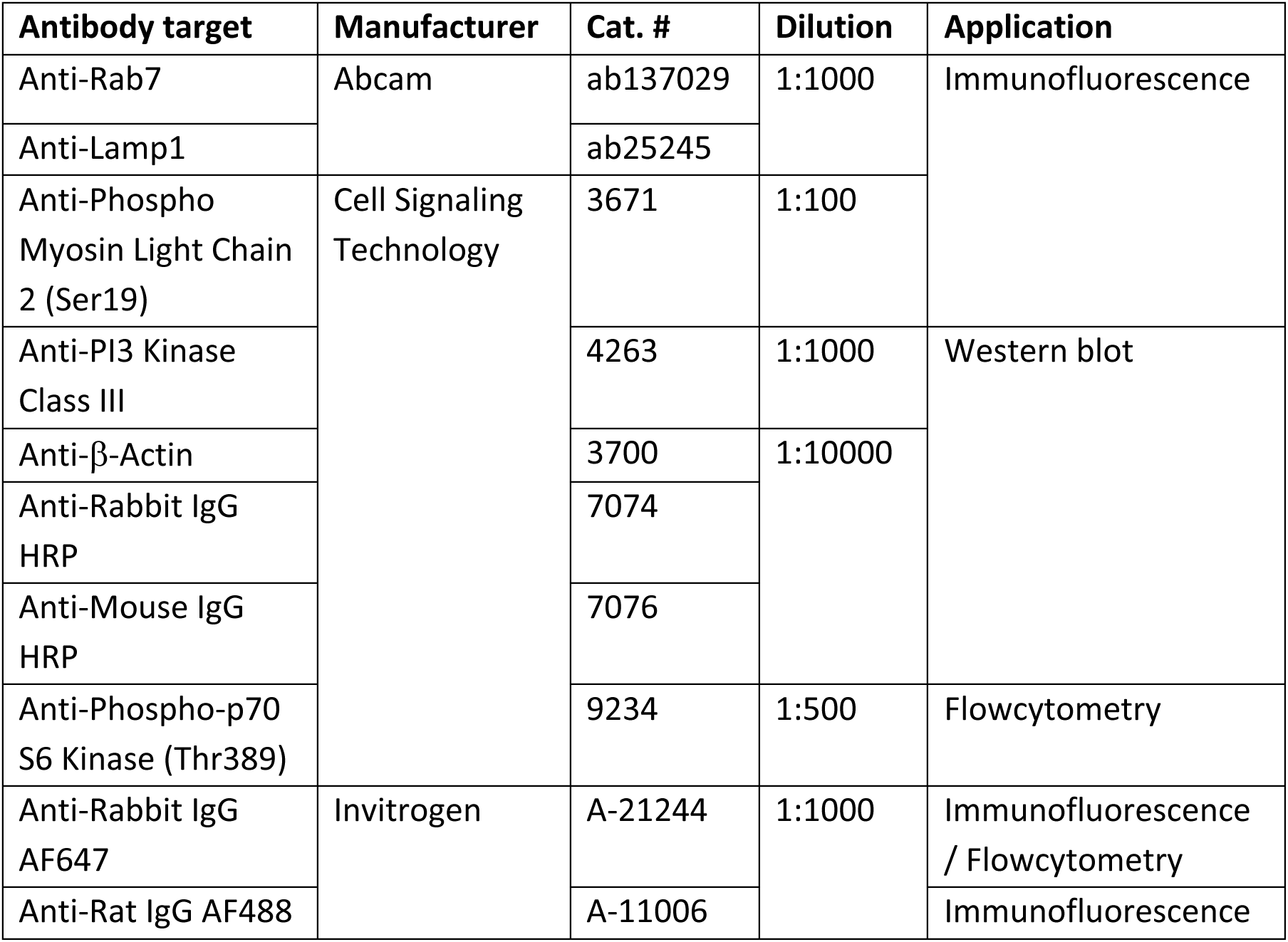

**Table 4.**

| Name | Target | Manufacturer | Cat. # | Concentration |
| --- | --- | --- | --- | --- |
| PQR620 | mTORC1/2 | Wymann Lab<br>(University of Basel) |  | 2 µM |
| VPS34-In1 | VPS34 | Selleckchem | S7980 | 2 µM |
| SAR405 | VPS34 | EMD Millipore | 533063 | 4 µM |
| Apilimod | PIKFYVE | MedChem Express | HY-14644A | 2 µM |
| (S(-4)′-nitro-Blebbistatin | Non-muscle myosin II ATPases | Cayman Chemical | 24171 | 25 µM |
| ML-7 | MLCK | MedChem Express | HY-15417 | 10 µM |
| Nocodazole | Microtubules | M1404 | M1404 | 1 µM |
| MK6-83 | TRPML1 | Sigma-Aldrich | SLM1509 | 5 µM |
| BAPTA-AM | Calcium | MedChem Express | HY-100545 | 20 µM |

### Immunoblotting

Human CD8^+^ T cell blasts were harvested, washed with ice-cold PBS, and then centrifuged at 500 x g for 5 min. Cells were lysed in RIPA buffer (ThermoFisher, Cat. # 89901) supplemented with protease inhibitors (Roche, Cat. # 05892970001) and phosphatase inhibitors (Roche, Cat. # 04906845001). Cell lysates were stored at -20 °C and cleared by centrifugation at 15000 g for 10 min at 4 °C prior to use. Protein concentrations were determined with a BCA Protein Assay Kit (ThermoFisher, Cat. # 23225). For each sample, 5 µg of protein was combined with 4x Laemmli buffer (Biorad, Cat. # 1610747) and 10 % 2-mercaptoethanol, and then denatured at 95 °C for 5 min. Protein samples were loaded into 4 - 20 % precast gradient gels (Biorad, Cat. # 1704158) alongside a page ruler (ThermoFisher, Cat. # 26620), and then subjected to electrophoresis for 5 min at 80 mV, followed by 55 min at 100 mV. The separated proteins were transferred to nitrocellulose membranes (Biorad, Cat. # 1704158) using a Trans-blot turbo system (Biorad) and blocked with 5 % BSA in TBST for 1 h at room temperature (RT). Membranes were sliced at the 70 kDa mark and incubated overnight at 4 °C with either anti-VPS34 or anti-Actin antibodies (**Table 3**). Following three washes with TBST, membranes were incubated for 1 h at RT with horseradish peroxidase (HRP) conjugated secondary antibodies (**Table 3**). Membranes were then washed, incubated for 5 min with chemiluminescent substrate (ThermoFisher, Cat. # 34580), and imaged on a gel doc system (Biorad).

### Treatment with Inhibitors / Agonists and Dipalmitoyl PI(3,5)P_2_ Supplementation

All inhibitor stock solutions were prepared in DMSO, with the same DMSO concentration applied for vehicle controls. Inhibitor concentrations and manufacturers are listed in **Table 4**. The compound PQR620 was synthesized in-house at the University of Basel’s Wymann Laboratory, following a published protocol.^24^ Prior to experiments, cells were treated with inhibitors or vehicle control for 2 hours. Exceptions were MK6-83 and Nocodazole, which were exclusively incorporated into the agarose gel for migration assays.

PI(3,5)P_2_-diC_16_ (Echelon Biosciences, Cat. # P-3516) was conjugated to fatty-acid-free BSA (Sigma-Aldrich, Cat. # A7030) by incubating at 37 °C for 1 h. Subsequently, phenol-red free RMPI1640, preheated to 37 °C, was added to the fatty-free BSA to attain a concentration of 100 mg / ml, yielding a final stock concentration of 500 µM. Aliquots of this stock were stored at -80 °C. As needed, the stock solution was diluted in complete medium for agarose gel preparation or cell treatments, as previously described.

### Staining of Live Cells with Fluorescent Dyes

For calcium imaging, naïve murine CD8^+^ T cells were stained with 2.5 µM Calbryte 520 AM (AAT Bioquest, Cat. # 20650) in complete medium at 37°C for 30 min. Subsequently, a 2 h drug treatment was carried out in the presence of 1µM LysoTracker Deep Red (Invitrogen, Cat. # L12492). Lysosensor Green DND-189 was added to the complete medium at a concentration of 1 µM and the cells were incubated at 37 °C for 2 h. Nuclear staining was performed using Hoechst 33342 (ThermoFisher, Cat. #R3705; one drop per two ml of medium) in parallel with the drug treatment.

### Electroporation of 2xFYVE-eGFP Plasmid into T cells

The pEGFP-2xFYVE plasmid, (a kind gift from Harald Stenmark; Addgene, Cat. # 140047) electroporation was performed using a 4D-Nucleofector (Lonza, Cat # AFF-1002B) with the P3 Nucleofector kit (Lonza, Cat. # V4SP-3096). Naïve murine CD8^+^ T cells (2 x 10^6^) were washed twice with PBS and resuspended in 20µl of P3 electroporation buffer containing 1 ng of the plasmid. The cell suspension was then added to an electroporation cuvette strip. Immediately after electroporation, 130 µl of prewarmed complete medium with 20 ng / ml IL-7 was added to the cuvette, and the cells were allowed to rest for 10 min at 37 °C. The cell suspension was mixed, centrifuged at 200 x g for 5 min, and resuspended in 1 ml of prewarmed complete medium with 20 ng / ml IL-7. The cells were prepared for imaging on the following day.

### Flow Cytometric Analysis of F-Actin Polymerization, Live / Dead Staining, S6K Phosphorylation, and CCL19 Uptake

For F-actin staining, 7.5 x 10^4^ naïve murine CD8^+^ T cells were treated with either a vehicle, VPS34In-1, or Apilimod and then stimulated for 1 min with 0.5 µg / ml CCL19. The cells were fixed directly in the medium with 4% PFA for 15 minutes, washed in PBS, and permeabilized with 0.2 % Triton-X in PBS for 15 minutes. F-actin was stained with AF-488 phalloidin (1 drop in 4 ml FACS buffer; Invitrogen, Cat. # R37110) for 30 minutes, washed thrice, and then acquired in FACS buffer (PBS & 5% BSA).

To assess the effect of VPS34 or PIKFYVE inhibition on murine CD8^+^ T cell blast viability, cells were treated with inhibitors for 6 hours and then stained with Zombie Aqua (BioLegend, Cat. # 423101) following the manufacturer’s instructions.

S6K phosphorylation was measured by intracellular staining in murine CD8^+^ T cells activated for 24 h with plate-bound anti-CD3 (coated at 5 µg / ml for 14 h at 4 °C in PBS; BioLegend, Cat. # 100331) and 1 µg / ml soluble anti-CD28 (BioLegend, Cat. # 1002116). Cells were stained for live/dead cells, fixed, and permeabilized for 15 min, followed by overnight staining with a primary anti-pS6K antibody at 4 °C and one hour of secondary antibody staining at RT (**Table 3**). This staining procedure was carried out with the Transcription Factor Staining Buffer Set (Invitrogen, Cat. # 00-5523-00).

CCL19 uptake in naïve CD8^+^ T cells was measured by adding 20 nM of DY-649P1-CCL19 to the cell culture medium for 0, 30, or 90 min. The cells were then subjected to live / dead staining and washed in FACS buffer. All samples were acquired with a CytoFlex (Beckman Coulter) and analyzed using Flowjo (BD).

### Preparation and Acquisition of Electron Microscopy Samples

A total of 1 x 10^7^ cells were fixed by adding a 2x concentrated EM fixative (comprising a final solution of 2.5 % glutaraldehyde (Sigma-Aldrich, Cat. # G5882) and 2.0 % paraformaldehyde (Science Services, Cat. # E15710) in 100 mM PIPES buffer (pH 7.2; Sigma-Aldrich, Cat. # P6757) to the cell culture medium (1:1) for 20 min at RT. After removing the fixative, it was replaced with a 1x fixative containing 0.01 % green malachite (Sigma-Aldrich, Cat. # 32745) for 2 h on ice. The cells were rinsed six times with PIPES buffer before being embedded in a 2 % low melt agarose (Roth, Cat. # 6351.1). Embedded cell pellets were washed five times for 3 min each in cacodylate buffer (0.15 M cacodylate with 4 mM CaCl2; Electron Microscopy Sciences, Cat. # 12300) before being post-fixed with osmium-ferricyanide (2 % OsO4, Electron Microscopy Sciences, Cat. # 19100; 1% K3Fe(III)(CN)6, Electron Microscopy Sciences, Cat. # 20150; in cacodylate buffer) for 30 minutes on ice. After five washes with ddH_2_O for 3 min each, cell pellets were immersed in a thiocarbohydrazide (TCH, Sigma-Aldrich, Cat. # 223220) solution for 30 min. Subsequently, cell pellets were treated with 1% OsO4 in water for 30 min at RT, washed five times, and immersed in 1 % uranyl acetate (Electron Microscopy Sciences, Cat. # 22400) overnight at 4 °C. On the following day, cells were washed and treated with Walton’s lead aspartate (0.66% lead nitrate in 0.4 % aspartic acid solution) for 1 h. After another wash, the cells underwent dehydration using an ethanol series (25 %, 50 %, 75 %, 90 %, and 2x 100 %) for 5 min each. The cell pellets were then infiltrated with Durcupan (Electron Microscopy Sciences, Cat. # 14040) using an ethanol-Durcupan gradient (20%, 50%, 70%, 90%, and 2x 100%) for 3 min each in the microwave. This was followed by a 30 min immersion in a 1:1 Ethanol:Durcupan resin and a subsequent overnight immersion in 100 % Durcupan solution. The next day, cells were immersed in freshly prepared resin for 1 h. Resin blocks were polymerized at 60 °C for 3 d. The polymerized samples were mounted on SEM stubs and imaged on either an FEI Helios NanoLab 650 (courtesy of NanoImaging Lab Basel; acquired pixel size: 8 x 8 x 16 nm) or a ZEISS Crossbeam 550 serial block face (courtesy of the Electron Microscopy Core Facility of EMBL Heidelberg; acquired pixel size: 8 x 8 x 40 nm). Image stacks were aligned using the SIFT module available on FIJI for analysis.^72^

### *In vivo* T cell homing assay

For adoptive T cell transfer, isolated murine CD8^+^ T cells were initially rested overnight and then divided into two groups. One group was labeled with 250 nM CellTrace Violet (CTV), while the other group was labeled with 1000 nM CellTrace Far Red (CTFR) (Invitrogen, Cat. # C34557 / C34564) for 15 minutes in PBS at 37 °C. Subsequently, the stained cells were separately treated with either 1:1000 DMSO or 2 µM Apilimod for 2 h at 37 °C, followed by washing. The cells were then combined in a 1:1 ratio in PBS. CD8^+^ T cells with wild-type VPS34 (isolated from CD8a-cre^+/-^, VPS34-Exon21^wt/wt^ animals) were labeled with CTV, while those with catalytic-dead VPS34 (CD8a-cre^+/-^, VPS34-Exon21^flox/flox^ animals) were labeled with CTFR, and both were mixed in a 1:1 ratio. Generation and characterization of the VPS34-Exon21 flox model was described previously.^38^ CD8a-cre and VPS34-Exon21-flox mice were bred and housed in a specific-pathogen-free facility at the University of Cambridge.

The labels were switched for each respective repetition experiment. For injection, a total of 3 x 10^6^ cells (1.5 x 10^6^ per condition) were administered into the tail vein of recipient mice. After 1 h spleens and lymph nodes (inguinal, axillary, and cervical) were harvested, and single-cell suspensions were obtained by passing the organs through a 70 µm cell strainer. Samples collected from spleens and organs were subjected to red blood cell lysis, followed by a single wash in FCS buffer and fixation with 4% PFA. The ratio of transferred cells was quantified using a CytoFlex (Beckman Coulter) or Aurora flow cytometer (Cytek). Subsequently, the homing index was calculated by determining the ratio of CTV-labeled cells to CTFR-labeled cells in comparison to the input ratio. The relative abundance of Apilimod-treated cells or cd-VPS34 T cells was normalized to their internal controls per recipient mouse.

### Sequence alignment, phylogenetic analysis and FYVE-domain retrieval

The amino acid sequences were accessed through the public domain (UniProt; **Table 5**).^73^ The Mega (Version 11.0.13) software was used to perform alignments with the MUSCLE algorithm and the phylogenetic analysis with the maximum likelihood method.^74^ The FYVE domains were extracted by searching the InterPro for the FYVE zinc finger domain (IPR000306) for the analyzed animals and filtering out duplicate protein names.^55^

**Table 5.**

| Organism | Gene / orthologue | Accession # |
| --- | --- | --- |
| *Arabidopsis thaliana* | VPS34 | P42339 |
|  | FAB1A / PIKfyve | Q0WUR5 |
| *Danio rerio* | PIKfyve | B2KTE3 |
|  | p110α | F1QAD7 |
|  | VPS34 | F1Q9F3 |
| *D. discoideum* | PI5K3 / PIKfyve | B0G126 |
|  | pikA / p110α | P54673 |
|  | pikE / *VPS34* | P54676 |
|  | Mucolipin / *TRPML1* | Q54Y0 |
| *Homo sapiens* | PIKfyve | Q9Y2I7 |
| | p110α | P42336 |
|  | VPS34 | Q8NEB9 |
| *Mus musculus* | Pikfyve / *PIKfyve* | Q9Z1T6 |
| | p110α | P42337 |
|  | VPS34 | Q6PF93 |
|  | TRM<sup>^</sup>MPL1 | Q99J21 |
| *Saccharomyces cerevisiae*<br>(*Saccharomyces cere.*) | VPS34 | P22543 |
|  | Fab1 / *PIKfyve* | P34756 |
| *Xenopus tropicalis* | PIKfyve | A0A6I8Q3Y5 |
| | p110α | B0BLW6 |
|  | VPS34 | F6ZM84 |

### Protein structure analysis and *in silico* docking

Protein structures for TRPML1 bound to PI(3,5)P_2_ (*Mus musculus; experimentally determined*; UniProt Entry: Q99J21, Structure identifier: 7sq7) and Mucolipin (*D. discoideum; Alphafold model*; UniProt Entry: Q54EY0, Structure identifier: AF-Q54EY0) were sourced through the public domain (UniProt, AlphaFold).^73,75,76^ Structures were aligned in PyMol and electrostatic potential was computed with the Adaptive Poisson-Boltzmann Solver (APBS) plug-in.^77^ The *in silico* docking was performed with high ambiguity-driven protein-protein docking (HADDOCK 2.4).^78,79^ The predicted mucolipin and PI(3,5)P_2_ structures were inputted and the PI(3,5)P_2_ binding AAs were added as interacting residues. The output model was then loaded into the PyMol and aligned with the TRPML1 structure.

---

**Fig. S1:**
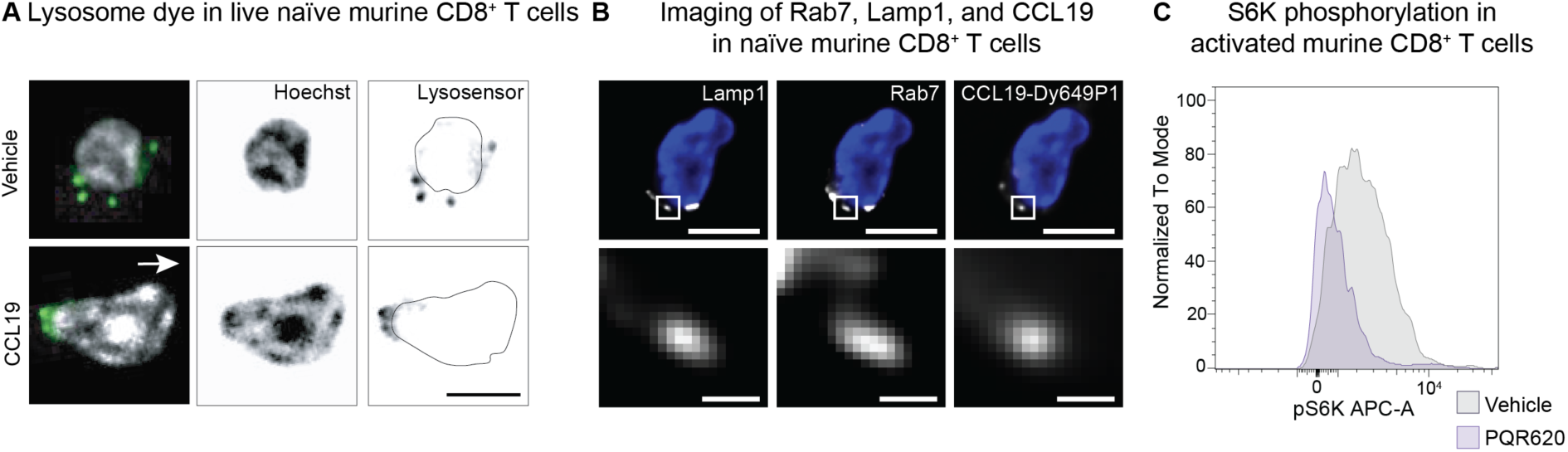
Endo-lysosomes decorated with VPS34–PIKfyve relocate to the uropod of polarized T cells |. **A**) Representative live cell images of lysosomes (Lysosensor dye, *right panels*), the nucleus (Hoechst dye, *middle panels*), and a merged images (*left panels*) of naïve murine CD8^+^ T cell treated with CCL19 or vehicle control. The arrow in the left panel indicates direction of migration and the black line in the right panels marks the nuclear outline. *Scale bar: 5 µm.* **B**) Representative fluorescence microscopy images of a naïve murine CD8^+^ T cell treated with CCL19-Dy-649P1 (*right panels*) and stained with Lamp1 (*left panels*) & Rab7 (*middle panels*). *Top panels*: Overlay with DAPI signal (blue), the box indicating the region of interest magnified in the *bottom panel*. *Scale bar: 5 µm (top panels), 500 nm (bottom panels).* **C**) Representative histogram (flow cytometry) of pS6K expression among murine CD8^+^ T cells activated for 24 h with CD3 and CD28 targeting antibodies and treated with the mTOR inhibitor PQR620 (purple) vs. vehicle control (grey).

**Fig. S2:**
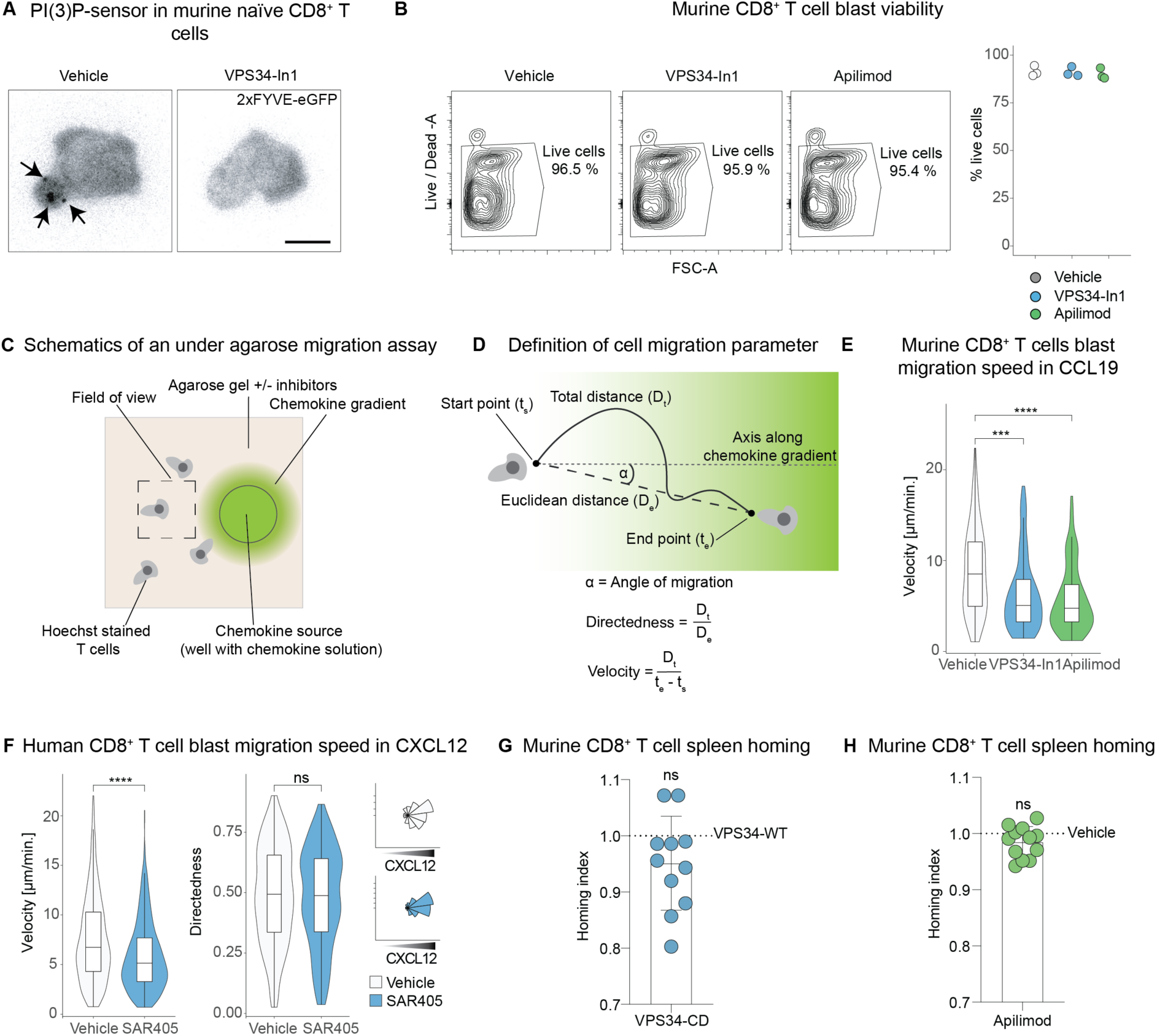
T cell velocity, but not directedness, is controlled by VPS34 and PIKfyve |. **A**) Naïve murine CD8^+^ T cells transfected with the 2xFYVE-eGFP PI(3)P sensor, placed in a CCL19 gradient and treated with vehicle control or VPS34-In1. Arrows point at PI(3)P^+^ vesicles. *Scale bar: 5 µm.* **B**) *Left panels*: Representative live/dead staining plots (flow cytometry) of murine CD8^+^ T cell blasts exposed to vehicle control, VPS34-In1, or Apilimod for 6 h. *Right panel:* Percentage of live cells of three biological replicates. **C**) Schematic (top view) of a T cell under-agarose migration assay. **D**) Definitions of cell migration metrics used in this report. **E**) Migration speed of murine CD8^+^ T cell blasts in a CCL19 gradient, treated with vehicle control (n = 66), VPS34-In1 (n = 81), or Apilimod (n = 79). **F**) Migration speed of human CD8^+^ T cell blasts migrating in a CXCL12 gradient, treated with the VPS34 inhibitor SAR405 (n = 626) or vehicle control (n = 755). **G**) Spleen homing index (normalized to each internal control) of murine CD8^+^ T cells expressing the wild-type or catalytic dead (CD) VPS34, and **H**) spleen homing index (normalized to each internal control) of vehicle or Apilimod-treated cells. *ns = p > 0.05, *** p < 0.001, **** p < 0.0001; Wilcoxon-test for panels E-H*.

**Fig. S3:**
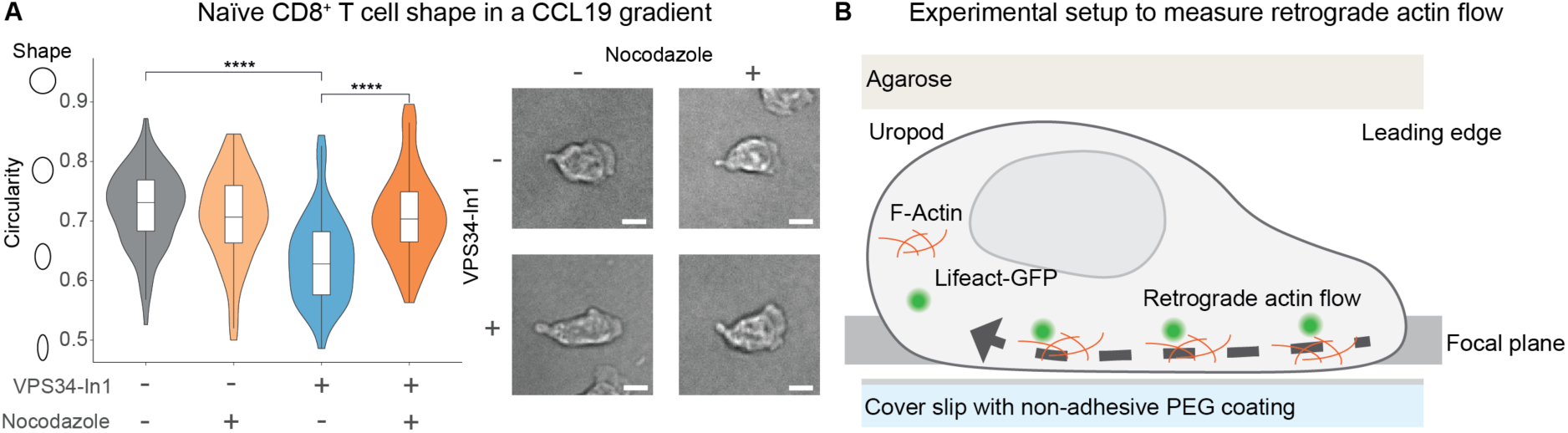
VPS34 and PIKfyve regulate retrograde actin flow and myosin IIA activation |. **A**) *Left panel:* Circularity of naïve CD8^+^ T cells placed in a CCL19 gradient treated with vehicle control (n = 81), Nocodazole (n = 70), VPS34-In1 (n = 85), and VPS34-In1 & Nocodazole (n = 56). *Right panels:* representative bright field images. *Scale bar: 5 µm. **** p < 0.0001, ART ANOVA.* **B)** Experimental setup for assessing retrograde actin flow in murine T cell blasts expressing the Lifeact-GFP reporter (to visualize F-actin dynamics).

**Fig. S4:**
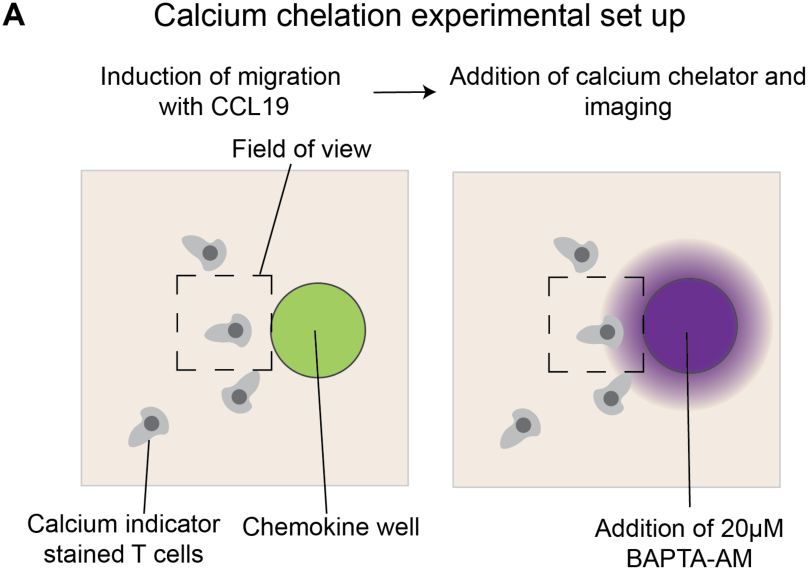
VPS34 and PIKfyve regulate migration velocity of T cells via lysosomal Ca^2+^|. **A**) Experimental setup of calcium chelation experiments. Naïve murine CD8^+^ T cells, stained with a calcium indicator (Calbryte dye), were treated with either VPS34-In1 or Apilimod and allowed to migrate in a CCL19 gradient (*left panel*). The chemokine solution in the well was then replaced with medium containing BAPTA-AM (a membrane-diffusible calcium chelator), and the calcium indicator signal in T cells was imaged over time (*right panel*).

**Fig. S5:**
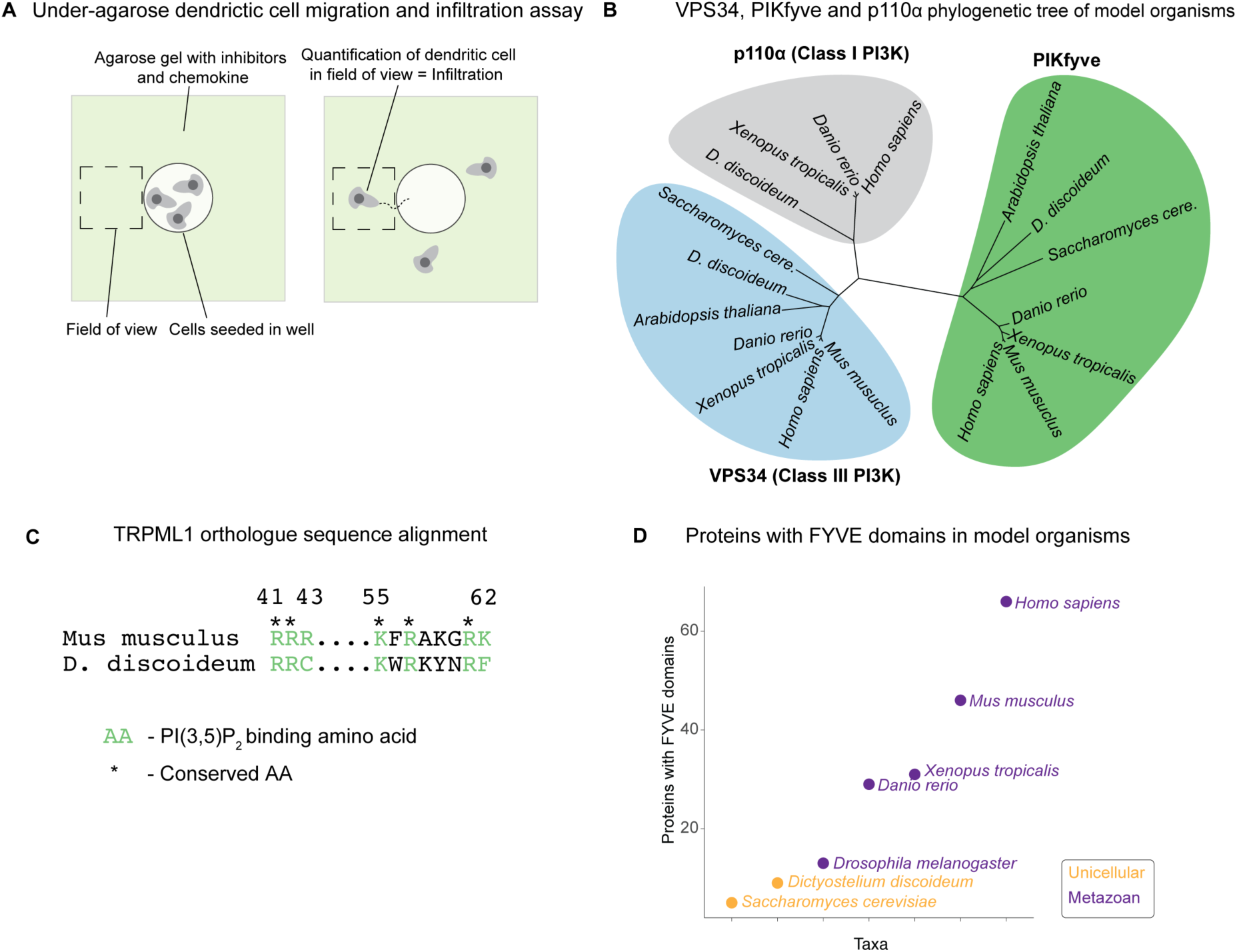
Regulation of amoeboid migration is a conserved function of the VPS34–PIKfyve pathway |. **A**) Schematics of the migration and infiltration assay of DCs: Cells were added to the well in the agarose (containing inhibitors and CCL19, *left panel*). DCs crawled under the gel (= infiltration) and the number of cells in the field of view was quantified (*right panel*). **B**) Phylogenetic analysis of the amino acid (AA) sequences of VPS34, PIKFYVE, and p110α (as an example for Class I PI3K). **C**) Alignment of the PI(3,5)P_2_-binding protein region from *Mus musculus* TRPML1 and the *Dicty* orthologue. PI(3,5)P_2_-binding AAs are highlighted in green, conserved AAs indicated by an asterisk. **D**) Number of proteins with a FYVE domain across the listed species. Orange indicates non-metazoan, purple metazoan organism.

